# Concordant yet unique neutral and adaptive genomic responses to anthropogenically modified landscapes in *Rhinella horribilis*

**DOI:** 10.1101/2025.01.29.635472

**Authors:** Gerardo J. Soria-Ortiz, Leticia M. Ochoa-Ochoa, Juan P. Jaramillo-Correa, Íñigo Martínez-Solano, Ella Vázquez-Domínguez

## Abstract

Anthropized environments are significantly challenging for wild species. Rapid adaptation to such habitats is thus key for their long-term persistence. Deciphering the environmental factors associated with species tolerance to modified habitats is fundamental for understanding the genetic and connectivity patterns of individuals and the local adaptation of their populations. We studied the Giant Toad, *Rhinella horribilis*, from two landscapes with distinct levels of anthropogenic habitat modification, assessed their genomic diversity, structure and connectivity with ddRAD-seq genomic data, identified potential outlier loci and their relationship with environmental and physicochemical water variables, and evaluated if populations from the two study sites showed signals of parallel adaptation. Both concordant and unique patterns were found regarding landscape factors and genotype-environment associations related with the degree of anthropic modification between landscapes. Genomic structure and connectivity were significantly associated with the presence of temporary water bodies, low vegetation cover, high humidity, solar radiation, and temperature. Notably, we identified both shared and distinct outlier SNPs and annotated functional genes for the two landscapes. Genes were enriched for biological processes and metabolic pathways, which were in turn correlated with environmental and physicochemical water variables. Genes and metabolic pathways were associated mainly with embryonic development, sexual maturation and immune responses. Studies such as this one, in an often-disregarded species, illustrate how parallel and un-parallel adaptive landscape genomic patterns arise in the stressful conditions of anthropized habitats.

## Introduction

Adaptive and non-adaptive evolution is crucial for populations to respond to environmental change. Non-adaptive forces include mutation, genetic drift, and recombination, which are primarily random, whereas adaptive forces like natural selection depend on the fitness (relative or absolute) of individuals within populations [1,2]. Rapid adaptation to anthropized habitats is key for the long-term persistence of species [3–5]. The global impact of human actions on ecosystems is rapidly altering their abiotic and biotic characteristics. Some direct effects include changes in the structure and configuration of the landscape matrix (e.g., vegetation cover, land uses, soil types) and in local microclimatic conditions (e.g., temperature, solar radiation, humidity), and the pollution of soils and water bodies derived from agriculture, livestock and industrial activities [6–8]. These modifications have negative effects on the survival and abundance of many species of flora and fauna and constrain the movement of individuals among remaining habitat fragments or patches. Reduction in individual dispersal, and the consequent loss of gene flow, usually promotes genetic drift and differentiation among populations, with detrimental consequences for the long-term maintenance of genetic diversity [9,10]. Nevertheless, some species can persist, and even thrive, in modified areas, facilitated by specific adaptations in their life history, ecology and functional traits that counteract the negative consequences of isolation [3,11–13]. Yet, these selected variants can be lost through the homogenizing effect of gene flow from neighboring populations. Hence, jointly characterizing patterns of functional connectivity and deciphering the environmental factors associated with species tolerance and/or facilitation in modified habitats is fundamental to understand the genetic basis of local adaptation to anthropization in wild populations [14–16].

Functional connectivity refers to the degree to which organisms effectively disperse across a landscape, which depends on the species life history traits and the structure and composition of the landscape [16,17]. In amphibians, some landscape factors negatively impact functional connectivity, including the loss of vegetation cover, the extent of impervious surfaces, road infrastructure, and other changes in land uses [18,19]. As an example, the Northern Two-lined Salamander (*Eurycea bislineata*) that has very low vagility, and the Spotted Salamander (*Ambystoma maculatum*) that is strictly aquatic, maintain connectivity among populations embedded in urban areas, which is facilitated by the presence of streams and riparian vegetation [18,20]. Although much less studied, different climatic [21] and physicochemical variables of water bodies used as breeding sites [8,18,19] are important in facilitating or limiting gene flow among amphibian populations. Identifying which environmental variables have higher leverage on patterns of functional connectivity is therefore essential to understanding whether local adaptations evolve in isolation or in the face of gene flow and what are the main extrinsic cues shaping selective pressures.

Microhabitats in anthropogenically modified sites are subject to rapid changes [6], which can exert strong selective pressures on amphibian populations in just a few generations [22].

Survival in such conditions has been associated with the evolution of novel local adaptations, as illustrated by species resilient to urbanized areas, including fish (*Fundulus heteroclitus*) [23], mammals (*Peromyscus leucopus*) [24], birds (*Parus major*) [25], reptiles (*Anolis cristatellus*) [26], and amphibians (*Rana [Lithobates] sylvatica*) [22]. Amphibians are considered to have considerable potential for rapid adaptation [27], for example to changes in temperature (Wood Frog *R. sylvatica*) [28], or in their ability to respond to *Batrachochytrium* fungal infections [29].

However, their adaptations to anthropogenic habitats are just starting to be investigated; in a recent example, Homola et al. [22] identified signals of selection in genes associated with salinity and light pollution in *R. sylvatica* from urban (disturbed) vs. rural (undisturbed) sites.

One of the most successful (common and abundant) amphibian species across different environments in the Neotropics is the Giant Toad *Rhinella horribilis*, which is naturally distributed along the Pacific and Gulf of Mexico coasts in Mexico, and through Central America to northern South America. This species tolerates a broad range of climatic conditions and can breed in water bodies with suboptimal characteristics like excess of potassium and sodium ions (i.e., detrimental conditions) [30,31]. Its medium-large body size, terrestrial habits, ability to reproduce in both shallow lotic (stream) and lentic (temporary) water bodies, high dispersal capacity and toxicity are some of the traits that make *R. horribilis* a resilient, successful competitor [32,33]. No population genomics studies have been performed with *R. horribilis*, despite its phylogenetic, ecological, and life history traits that make it an excellent system to evaluate functional connectivity patterns and explore potential adaptation associated with environmental conditions in modified habitats.

In this study, we aimed to: (1) characterize the genetic diversity and structure of *R. horribilis* populations in two landscapes with distinct levels of habitat modification; (2) determine landscape variables (climatic, vegetation, land use) associated with functional connectivity; (3) identify outlier loci potentially associated with adaptation to modified habitats, assessing their relationship with landscape variables; and (4) test for signs of parallel adaptation (i.e., similar outlier loci and/or metabolic pathways) among populations of the two landscapes. Based on *R. horribilis* life history traits, we predict high gene flow (functional connectivity) among populations; also, positive genotype-environment associations reflecting tolerance to detrimental environmental and water conditions and potentially facilitating local adaptation to survive in human-modified habitats.

## Materials and methods

### Field sampling, environmental data and landscape features

We worked in two landscapes in the Sierra Madre del Sur, along the Pacific slope in the state of Oaxaca, southern Mexico. The landscapes are separated by ca. 70 km; each has a different configuration, amount of vegetation cover, and land uses (Fig 1). Landscape 1 (P1O) has lower vegetation cover, higher fragmentation and more cultivated, pastures and livestock areas than the second landscape (P2O). Sampling was performed during the rainy season (May to September 2019). We chose eight and seven sampling sites in P1O and P2O, respectively, located on either side of a main road or rural town (urbanized area) (Fig 1) and separated by a minimum of 1.5 km and a maximum of 23 km (P1O) and 15 km (P2O) distance. This sampling scheme allowed us to test for isolation by barrier, since roads and highly modified habitats are known to represent significant barriers for the connectivity of amphibian populations [18,34]. We set two 50x2 m (100 m^2^) transects at each sampling site, which were sampled once each for 1.5-2 hours. We collected toe clips from adult (post-metamorphic) *Rhinella horribilis* individuals; tissue samples were stored in eppendorf tubes with 99% ethanol; all individuals were released at their sampling location. Fieldwork, sampling and animal care were performed in strict adherence to the guidelines for working with amphibians and reptiles [35], approved and following the guidelines of the Ethics Committee Facultad de Ciencias-UNAM (project PN 2271), and with the appropriate scientific collecting permit from Secretaría del Medio Ambiente y Recursos Naturales (SEMARNAT; 09/K4-1472/09/18 to LMOO). We followed AmphibiaWeb (https://amphibiaweb.org/) for taxonomy nomenclature.

**Fig. 1.**
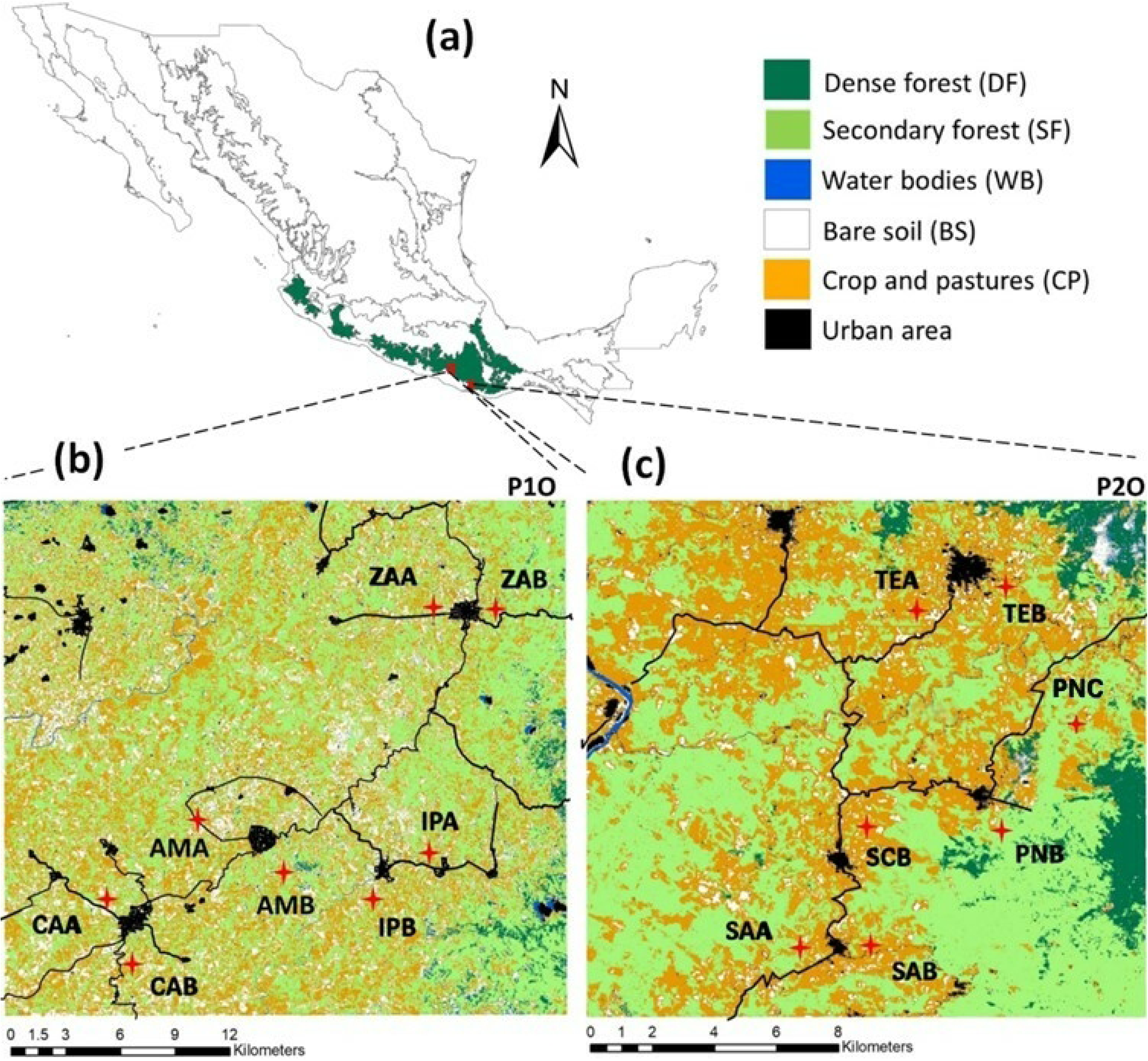
Landscape structure and sampling site location for *Rhinella horribilis* within the state of Oaxaca, southern Mexico. (**a**) Sierra Madre del Sur (green outline), (**b**) Landscape 1 (P1O), (**c**) landscape 2 (P2O). The sampling sites are indicated with red crosses; note they are located on both sides of a road and/or urban town. (**a**) and (**b**) have different scales due to distinct extent of study area. See Table S3 for sampling sites names.

To characterize the local environment of each landscape, we obtained *in situ* data on climatic and water body variables and of landscape features. For the first, we used a Kestrel 3000 climate meter (Kestrel® Instruments) to take measures of ambient temperature (AT) and relative humidity (RH) along the transects at each sampling site. We also obtained data for the solar radiation (SR) and evapotranspiration (EVA) variables from WorldClim 2.1, at a resolution of 1 km. For water variables, we used a Hanna HI9829 multiparameter meter (HANNA® Instruments) to measure six physicochemical water properties in all the ponds and streams where toads were sampled: pH (pH), water temperature (WT), oxygen availability (OA), and particles per million (ppm) of potassium (K), nitrates (NO), and sodium (Na). For the landscape variables, we used the ‘Mexican elevational continuum surfaces’ v3.0 (resolution of 15 m, available at https://www.inegi.org.mx/app/geo2/elevacionesmex/) to generate the elevation variable (ELV); and extracted information on the main and secondary roads from the National Road Network (http://rnc.imt.mx, downloaded in October 2023) at 10 m resolution; and of rivers and streams (10 m) from the INEGI national hydrology database (https://www.inegi.org.mx, downloaded in October 2023). Finally, we downloaded satellite images (May 2019) from the Landviewer platform (Sentinel-2 L2A) at a 10 m resolution, and estimated indices of normalized vegetation (NDVI), normalized humidity (NDMI) and bare soil (BSI) with QGIS v3.22. We used infrared visualization to identify temporary water bodies in the satellite images (see [36]); briefly, we did a kernel density analysis with a 500 m search radius from unique points representing potential water bodies (pixels) and temporal streams (lines). Based on this information, we identified areas with a high probability of presence of temporary water bodies (in the rainy season). For all variables, we built environmental layers with QGIS v3.22 [37], which were restricted to the study areas and reclassified to a pixel size of 30x30 m.

### DNA extraction, sequencing, and bioinformatics processing

We extracted genomic DNA from 190 *R. horribilis* individual tissues with the DNeasy Blood and Tissue Kit (Qiagen, Valencia, CA, USA), following the manufacturer’s protocol. We confirmed DNA quality and quantity on 1% agarose gels using GelRed® (Biotium, Fremont, CA) and a Qubit^TM^ fluorometer (Invitrogen, Carlsbad, CA). We randomly selected 10% of extractions to perform *in silico* digestions with SimRAD v.0.96 [38] in R [39] and identified, using 5% of the genome of *Rhinella marina* (Ensembl: GCA_900303285.1; version: gi:1370235108), the best cutting restriction enzymes (*Pst*I and *Bfa*I). Library preparation and sequencing were performed at the University of Wisconsin Biotechnology Center using the paired-end ddRAD protocol [40] and the selected restriction enzymes. Samples were sequenced on an Illumina NovaSeq line to obtain 150-bp paired reads.

The bioinformatic processing comprised the following steps: demultiplexing raw reads with the function *process_radtags* in Stacks v.2.62 [41]; quality filtering with FastQC v.0.11.9 [42]; removing adapters and discarding reads with Phred quality score <20, and trimming reads to a minimum length of 100 bp with Trimmomatic v.0.36 [43]. We aligned the resulting reads to the *R. marina* reference genome using the function *BWA mem* in BWA v.0.7.1 [44]. Calling of single nucleotide polymorphisms (SNPs) was done with the *ref_map.pl* pipeline and the *populations* module in Stacks, using the parameters p=15, r=0.7.

The resulting SNPs dataset was separated by landscape (P1O and P2O) and additional quality filtering was done separately for each one with VCFtools v.0.1.13 [45]. Considering our main objectives, we built two SNPs datasets for each landscape; namely, to characterize genetic diversity and structure and to evaluate landscape genomics of *R. horribilis* populations in each landscape (neutral processes), a stringent filtering [46] was performed for the first dataset (‘neutral data’): minimum depth of 10 samples to recognize a SNP as true, maximum allowed amount of missing data = 20%, and minimum allele frequency (maf) = 0.1. Missing data by locus and by individual >20% were removed, only biallelic loci were retained, and loci out of Hardy-Weinberg equilibrium (*p* >0.05) were eliminated. For the detection of outlier SNPs and genomic regions potentially under selection, a second less stringent dataset (‘full data’) was used, with parameters r=0.5, maf=0.05 and HWE=0.001 [46].

### Genetic structure and gene flow patterns

We calculated observed (*H_o_*) and expected (*H_e_*) heterozygosity and *F_IS_* values in each landscape with *populations* in Stacks. Genetic structure analyses were conducted on each landscape separately, using the neutral data and an array of complementary methods. We first applied two approaches that do not assume an underlying population genetic model, a principal component analysis (PCA) that is not affected by unequal sample sizes [47], and a Discriminant Analysis of Principal Components (DAPC), that uses the *a*-score method to determine the proportion of successful reassignment by individual as a function of the number of retained PCs, which maximizes the differentiation between populations. We ran PCA with the *glpca* function in adegenet v.2.1.3 in R [48], retaining 50 and 26 factors for P1O and P2O, respectively. DAPC was run with *dapc* in adegenet; individuals were grouped according to the eight and seven sampling sites in P1O and P2O, respectively, and the number of retained PCs (P1O=40, P2O=9; S1 Fig) was obtained with *xvalDapc* in poppr v.2.9.3 [49]. Additionally, we ran sPCA with *spca* in adegenet, a spatial model that is well suited to identify cryptic genetic structure, as it combines the spatial autocorrelation of sampling sites and the genetic variation within sites to maximize the effects of global and local structure [50]. Finally, we used two methods that are based on sparse non-negative matrix factorization, snmf [51] and TESS [52]. Both tests compute regularized least-square estimates of admixture proportions to estimate individual ancestry coefficients; TESS further includes the spatial information (geographic coordinates) of each sampling site. We tested from *K*=1 to the maximum number of populations in each landscape (*K*=8 in P1O and *K*=7 in P2O), with 20 replicate runs per *K-value*, using the function *snmf* in LEA v.3.2.0 [51] and *TESS3* in Tess3r v.1.1.0 [52], both implemented in R.

### Landscape genetics analyses

We performed correlation analyses, based on the layers we built for the different environmental variables per landscape, using function *cor* with stats in R. The normalized vegetation (NDVI), normalized humidity (NDMI) and bare soil (BSI) indices had a correlation coefficient ≥ 0.8 in both landscapes, while evapotranspiration (EVA), solar radiation (SR) and elevation (ELV) were also significantly correlated in P2O. Thus, we retained NDMI for the two landscapes and SR for P2O, to perform the connectivity analyses. We estimated pairwise genetic distance (*Fst*) between sampling sites within each landscape using *pairwise.neifst* with hierfstat v.0.5-11 in R [53], and the geographic (Euclidean) distances with fossil 0.4.0 [54] in R, with which we performed a Mantel test to assess isolation by distance. Next, we evaluated the influence of landscape features on gene flow and established connectivity hypotheses (predictions to be tested; see Table S1), considering the biology and ecology of *R. horribilis*. Specifically, we selected environmental variables that can potentially promote or impair individual movement and, ultimately, have a role on adaptative processes.

Considering that major roads and rivers are known to limit amphibian connectivity (e.g., [18]), we hypothesized that these features would function as barriers for *R. horribilis* movement and gene flow (Table S1). We assessed the relationship between genetic distance and both main roads and high-flow rivers that divide sampling sites within each landscape using two methods. We first performed a distance-based redundancy analysis (db-RDA) [55] that carries out a constrained ordination using non-Euclidian distance measures. We created a series of dummy variables [0, 1] to indicate that a sampling site was on one side of the road/river or the other, and used *Fst* values to perform a principal coordinate analysis (PCoA). The resulting eingenvectors were then used as a response matrix in a RDA [56], with which we tested road/river ‘barrier-effects’. We also applied a multiple regression on distance matrices (MRDM) incorporating multiple response variables [57] to test multiple barrier scenarios (see S2 Fig). To do so, we built resistance surfaces for each barrier scenario, assigning a value of 1000 to roads and rivers and 1 to the rest of the landscape. We obtained least-cost distance matrices for each scenario (response variables) as estimated with *costDistance* in gdistance v.1.6.4 in R [58], and the *Fst* matrix was used as exploratory variable. MRDM was run with *MRM* in ecodist v.2.0.9 in R [59], with 10,000 permutations.

Although amphibian distribution is in general constrained by humidity [60], *R. horribilis* can be present across open areas and can hence tolerate harsh temperature and radiation conditions [31,32]. Likewise, *R. horribilis* is often found in areas with low vegetation cover [31], but rarely near/at urban cities, while bufonids in general prefer more humid habitats [61,62].

Accordingly, we hypothesized that extremely high temperature, evapotranspiration and solar radiation (i.e., detrimental conditions) will limit *R. horribilis* movement in areas with no vegetation cover, while it will be facilitated in areas with higher vegetation cover, humid soils and lower elevation; also, that impervious surfaces are expected to function as barriers to movement (Table S1). Lastly, given that water bodies are crucial for amphibian reproduction and can function as stepping-stones, facilitating movement of individuals and connecting breeding sites [7,36,63], we predicted that areas with a higher probability for the presence of temporary water bodies (ponds and streams) will promote connectivity.

The connectivity models were performed with the optimization framework developed by Peterman [64], to determine the resistance values of our landscape variables surfaces with ResistanceGA [64,65]. This method uses a genetic algorithm (GA) [66] that adaptively explores the parameter space of monomolecular and Ricker functions [67] to transform continuous surfaces into resistance surfaces without prior assumptions. The monomolecular [y = *r* (1-exp^-bx^)] and Ricker [y = *r* exp^-bx^] are two exponential-based functions used for ecological modeling, which differ in the curve shape of the relationship they are modeling. The curve shape is mainly determined by shape (x) and magnitude (b) parameters, which result in a saturating exponential (growth or decay) curve for the monomolecular function, and a hump-shaped curve (skewed to right or left) for the Ricker function [67]. During the optimization process, the genetic algorithm searches all possible combinations of these parameters for transforming resistance surfaces, denoted by “r” in the monomolecular and Ricker equations [64,65]. The method also uses a maximum-likelihood population effects mixed models (MLPE) to evaluate the fitness of each surface, considering the non-independence inherent to pairwise distance matrices. In the process, the GA seeks to maximize the relationship between pairwise genetic distances (*F_ST_*; response variable) and pairwise landscape distances (predictor variables), including the Euclidean distance model. First, surfaces were individually optimized using the *commuteDistance* function with gdistance in R [58], with three independent runs to verify convergence of parameters. Support for the optimized resistance surfaces was assessed with AICc (Akaike’s information criterion corrected for small/finite sample size) [68], and a bootstrap resampling of the data to evaluate the robustness of the models. To do so, we randomly selected 75% of the samples without replacement and fitted each surface to each sample subset; the average rank, average *R^2^* (proportion of the variance explained by the fixed factors), and proportion (percentage) in which a surface was chosen as the best model were estimated with *Resist.boot* function [69], with 10,000 iterations. Finally, considering the best-supported individual models (Table 1) and our connectivity hypotheses, we built five composite surfaces (aquatic, environmental, global, biological-1 (facilitating) and biological-2 (limiting) models; see Table S2) to run a multivariate optimization approach. Optimization of composite surface models was conducted twice to ensure convergence [64]. Bootstrap model selection was performed again with the same previous parameters to obtain the average rank, average *R^2^*, and the selection percentage of both univariate and multivariate surfaces.

**Table 1.** Individual generalized linear mixed models for *Rhinella horribilis* in two landscapes (P1O and P2O) in Oaxaca, southern Mexico. Best-supported models are indicated by the highest ‘top model (%)’, k: number of parameters fit in each model plus the intercept, AICc: Akaike information criterion. The average rank and average *R^2^* (value of the fitted model; %) are shown.

| Landscape 1 (P1O) |  |  |  |  |  |  |
| --- | --- | --- | --- | --- | --- | --- |
| Surface | k | AICc | AICc | Average rank | Average $R^2$ (%) | Top mode |

|  |  |  |  |  |  | I<br>(%) |
| --- | --- | --- | --- | --- | --- | --- |
| Temporary water bodies<br>(TBW) | 4 | -116.55 | 0 | 1.5328 | 80.86 | 85.62 |
| Distance (D) | 2 | -106.96 | -9.59 | 4.3226 | 44.21 | 0 |
| Temporary streams (TS) | 4 | -106.30 | -10.25 | 5.9758 | 54.52 | 7.19 |
| Evapotranspiration (EVA) | 4 | -105.20 | -11.35 | 7.4184 | 48.39 | 0 |
| Solar radiation (SR) | 4 | -104.71 | -11.84 | 8.3501 | 44.68 | 3.81 |
| Normalized humidity (NDMI) | 4 | -104.26 | -12.29 | 8.4249 | 51.53 | 0 |
| Ambient temperature (AT) | 4 | -104.25 | -12.30 | 9.1776 | 44.73 | 3.38 |
| Bare soil (BSI) | 4 | -104.09 | -12.46 | 9.7481 | 50.74 | 0 |
| Relative humidity (RH) | 4 | -104.14 | -12.41 | 9.4443 | 46.73 | 0 |
| Elevation (ELV) | 4 | -103.21 | -13.34 | 10.6832 | 42.18 | 0 |
| Normalized vegetation<br>(NDVI) | 4 | -103.14 | -13.41 | 11.773 | 44.80 | 0 |
| **Landscape 2 (P2O)** |  |  |  |  |  |  |
| Distance (D) | 2 | -67.25 | -0.87 | 2.8877 | 40.95 | 75.97 |
| Ambient temperature (AT) | 4 | -68.27 | 0 | 2.3625 | 61.90 | 13.48 |
| Solar radiation (SR) | 4 | -65.88 | -2.19 | 4.2437 | 58.13 | 7.17 |
| Temporary water bodies<br>(TWB) | 4 | -65.23 | -3.04 | 6.5282 | 44.35 | 1.61 |
| Relative humidity (RH) | 4 | -65.03 | -3.14 | 5.4863 | 44.98 | 1.74 |
| Normalized humidity (NDMI) | 4 | -64.53 | -3.65 | 6.4141 | 43.92 | 0 |
| Evapotranspiration (EVA) | 4 | -64.32 | -3.95 | 5.8418 | 43.03 | 0 |
| Bare soil (BSI) | 4 | -64.17 | -4.10 | 7.4666 | 41.82 | 0 |
| Elevation (ELV) | 4 | -63.87 | -4.40 | 7.1094 | 41.62 | 0 |
| Temporary streams (TS) | 4 | -62.96 | -5.14 | 8.6201 | 41.11 | 0 |
| Normalized vegetation<br>(NDVI) | 4 | -63.25 | -5.02 | 8.5716 | 40.95 | 0 |

### Detection of outlier SNPs and genomic regions potentially under selection

We identified candidate loci with an outlier method using the function *pcadapt* in PCAdapt v.4.3.5 in R [70], which detects outlier loci based on principal component analysis (PCoA) and assumes that markers excessively related to population structure are candidates for potential adaptation; it does not require grouping individuals into populations and can handle admixed individuals. PCAdapt uses the genomic inflation factor (GIF) to correct for inflation of the test score at each locus, which can be due to population structure or other confounding factors. We ran this analysis with default parameters for *K*=3 latent factors (according to our genetic structure results) and assessed significance levels with a false discovery rate (FDR) procedure [71]; we retained all SNPs with FDR <0.01.

To select uncorrelated variables for GEA analyses, we performed correlation tests (≥ 0.8) of the climatic and water body variables measured *in situ* at each landscape, which resulted in the removal of variables pH and NO for P1O and pH and AT for P2O; then we performed a PCA comparing the final variables between the two landscapes. We applied two GEA approaches, first a redundancy analysis (RDA) [72] that identifies covarying loci associated with multivariate predictors (i.e., landscape variables), using *rda* in vegan v.2.6-4 in R [73]. Environmental variables were used as explanatory and loci as response variables. We assessed the number of canonical axes with the function *anova.cca,* and only significant axes were used to find outlier loci. Loci deviating more than 3 standard deviations from the mean (*p* ≈ 0.001) were retained as candidates and assigned to the variable with the highest correlation coefficient. Secondly, we applied latent factor mixed model (LFMM) with the function *lfmm_ridge* in lfmm v.1.1 [74] available in LEA in R [75]. LFMM detects correlations between environmental variables and genotypic variation to reveal outliers while controlling for population structure; it is efficient for dealing with false discovery rates, sampling design limitations, and spatially autocorrelated populations (i.e., IBD) [76,77]. We ran these analyses with *K*=3 (according to the genetic structure results) and 1,000 iterations.

### SNP annotation, enrichment analysis, environmental association and assessment of parallel evolution

We identified the contigs that contained the candidate SNPs and annotated them through searches in the *R. marina* (NCBI Taxonomy ID: 8386) and *Xenopus tropicalis* (NCBI Taxonomy ID: 8364) reference genomes, using the BLASTn tool in the NCBI nucleotide database (National Center for Biotechnology Information, https://blast.ncbi.nlm.nih.gov/Blast.cgi; accessed on January 2024) and the Xenbase (https://www.xenbase.org/xenbase/; accessed on January 2024), with an E-value cut-off of 0.05 and similarity >80%, and associated the gene names to Xenbase_ID identifier using DAVID at NCBI (https://david.ncifcrf.gov; accessed on January 2024). All annotated candidates were used to perform enrichment analyses in the ShinyGo v.8.0 platform (http://bioinformatics.sdstate.edu/go//) [78], with the *X. tropicalis* as reference, to identify overrepresented GO ontology terms [79] and KEGG pathways [80].

Given that our purpose was to test for signs of parallel adaptation (i.e., similar outlier loci and/or metabolic pathways) among populations of the two landscapes, we performed gene enrichment using those genes shared by the two landscapes (N=34; see Results). Additionally, we identified individual processes within landscapes, based on unique genes (P1O=228; P2O=106; see Results). Enriched GO terms and KEGG pathways were considered significant with an FDR ≤0.05. Next, we did a RDA using the SNPs associated with the 34 shared genes (explanatory variable) and the landscape variables of each landscape (response variable), to determine the relationship and direction between the environment and the genes identified with GO terms and KEGG pathways.

We aimed to identify if the potentially adaptive variation in the two landscapes responded similarly to the same landscape variables. To this end, we performed RDAs using different sets of SNPs (see S3 Fig for the testing scheme). First, we used those candidate loci identified uniquely in each landscape (see Results): (a) 609 SNPs in P1O (Set-609-P1O), (b) 237 SNPs in P2O (Set-237-P2O), while also (c) testing the 609 SNPs of P1O and (d) the 237 SNPs of P2O each in the other landscape (Set-609-P2O and Set-237-P1O, respectively). Additionally, the three methods consistently detected 14 outlier loci in P1O (see Results), so we tested these 14 in (e) their corresponding P1O landscape, and in (f) the other landscape P2O (Set-14-P1O, Set- 14-P2O). Next, to confirm the environmental genetic associations identified, we performed RDAs with the SNPs obtained from the ‘neutral data’ for each landscape (see Results): (g) 2391 neutral SNPs in P1O and (h) 3494 in P2O (SetN-2391-P1O and SetN-3494-P2O, respectively). To additionally test for an effect of the data set and of the sample size (number of SNPs), we randomly selected 609 and 237 neutral SNPs for each landscape (i) SetRN-609-P1O, (j) SetRN- 237-P1O, (k) SetRN-609-P2O and (l) SetRN-237-P2O, assessing the random-neutral loci on the other landscape as well (m, n, o, p; see S3 Fig; Table S3). Hence, we carried out a total of 16 tests, nine evaluating each landscape’s SNPs set with its landscape variables, and seven crossed tests, assessing the role of SNPs identified in one landscape in explaining environmental variation in the other landscape. Finally, to compare the overlap of the SNPs associated with the same climatic and physicochemical variables in the two landscapes with those under random expectations we used a Fisher’s exact test (*p* ≤0.05).

## Results

A total of 190 *R. horribilis* adults were sampled, 125 in landscape P1O and 65 in landscape P2O (Table S4). Genomic results yielded 4,356,669 polymorphic sites before filtering. The filtered ‘neutral data’ included 2391 SNPs for P1O and 3494 SNPs for P2O, with an average coverage of 25x. This data was used for the genetic diversity and structure analyses and for landscape genetics. The ‘full data’ set had 8753 and 13,544 SNPs, for P1O and P2O respectively (coverage 24x), which was used for the outlier detection.

### Genetic diversity and structure

Genetic diversity (average observed heterozygosity) was significantly higher in P2O (*Ho*=0.282) than P1O (*Ho*=0.266) (t = -5.9; *p* <0.001), while in all sampling sites, the observed heterozygosity was higher than the expected heterozygosity, and no significant *F_IS_* values were inferred (Table S4). Overall genetic differentiation (*F_ST_*) was higher in P1O than in P2O (average *F_ST_*=0.0151, P1O; *F_ST_*=0.0130, P2O). Within P1O, the site Santa María Zacatepec-B (ZAB) was most differentiated from the rest (*F_ST_* >0.0154; Table S5), while in P2O, the highest differentiation was observed between Santa Cruz Tepenixtlahuaca (TEA) and Santa Ana Tututepec (SAA) (*F_ST_*=0.0292).

Genetic structure results in P1O identified *K*=2-3 genetic groups; TESS, SNMF, and PCA separated sites from the north (ZAA and ZAB) and the center (AMA and AMB) of the study region from the remaining localities (CAA, CAB, IPA, IPB), which had varying levels of admixture with the two former groups (Fig 2a-c; S4a Fig). On the other hand, sPCA and DAPC showed that the northern (ZAA and ZAB) and southern (CAA and CAB) sites formed distinct genetic groups, with a third one formed by the central sites (Fig 2d,e), more in line with the spatial distribution of the sampled localities. Importantly, we corroborated that each sampling site did not form a distinct genetic group, implying that the roads and rivers separating each pair of sites do not impede gene flow (S4b-d Fig; *K*=3 in SNMF, *K*=8 in TESS). Patterns were similar for P2O (*K*=2- 3), where two genetic groups were identified, one in the northern portion (TEA and TEB) and one in the southern part of the study region (SAA and SAB); with the three remaining sites admixed with both groups (Fig 3a-c; S5a Fig). Again, sPCA and DAPC indicated that these central sites formed a different genetic group (Fig 3d,e). As for P1O, exploring higher values of *K* did not provide further genetic structure resolution (S5b,d Fig).

**Fig. 2.**
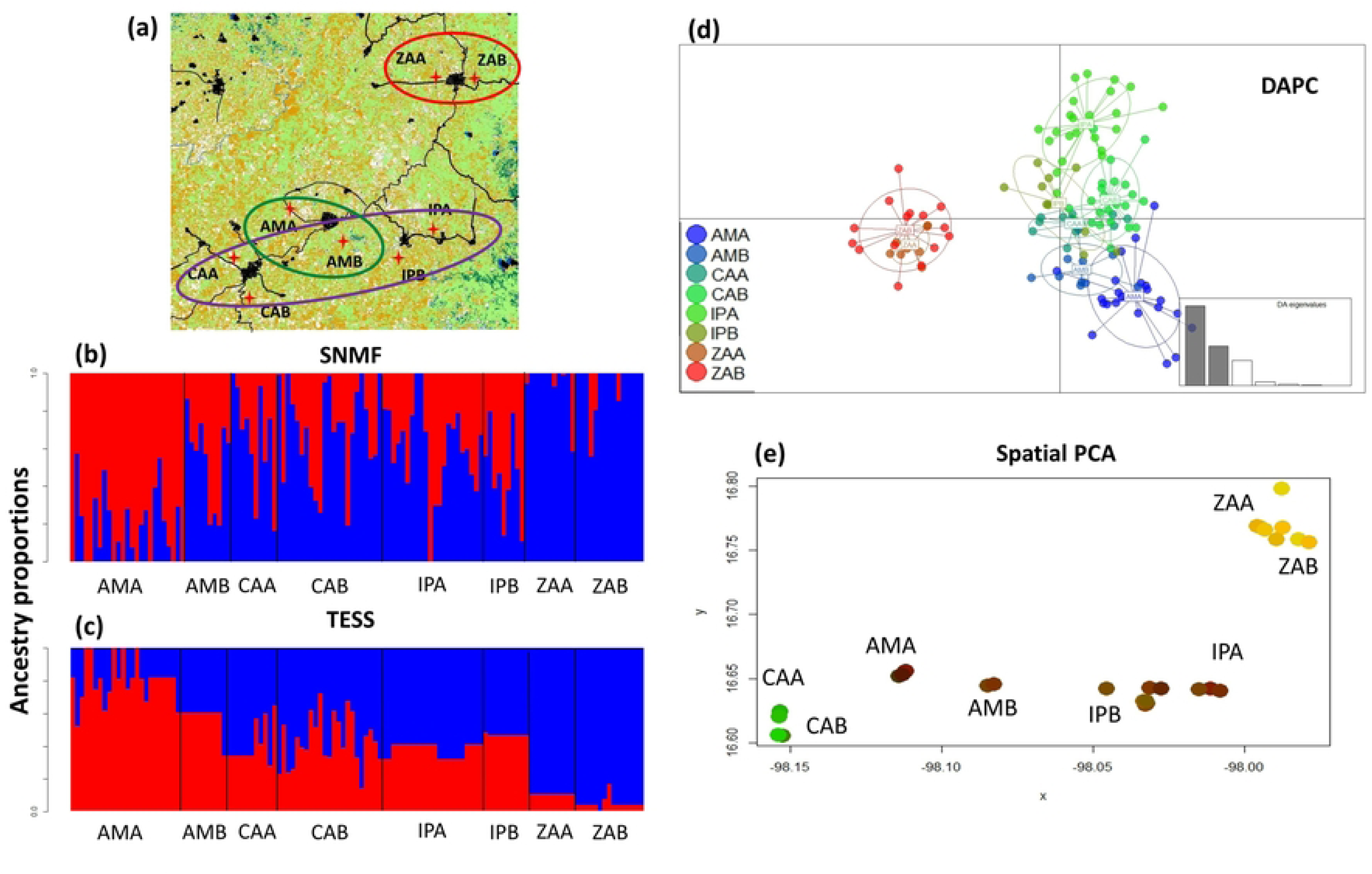
Genetic structure of *Rhinella horribilis* in landscape 1 (P1O) as estimated with different methods. (**a**) Sampling sites in Oaxaca, southern Mexico, (**b**) SNMF (*K*=2), (**c**) TESS (*K*=2), (**d**) DAPC, (**e**) sPCA.

**Fig 3.**
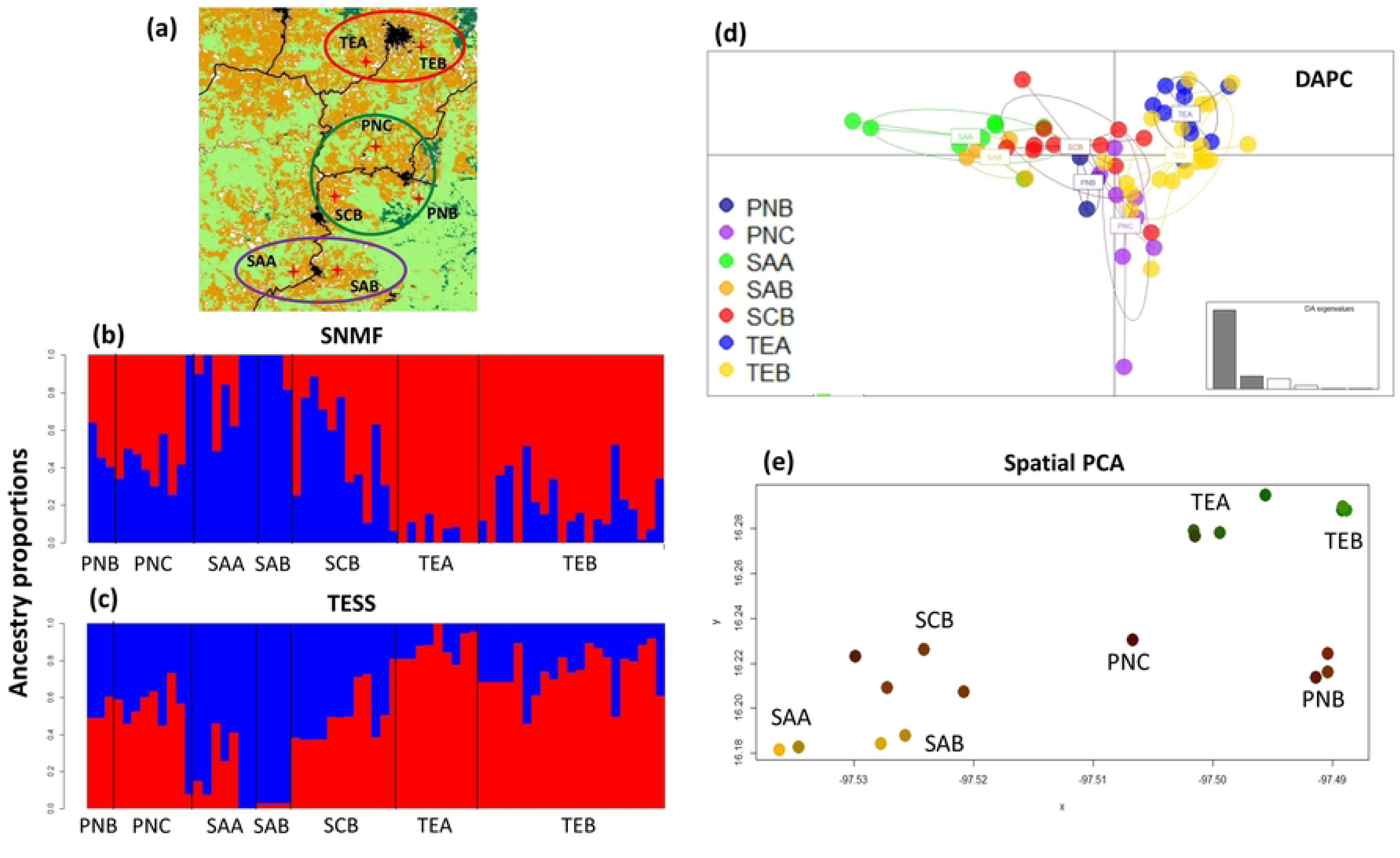
Genetic structure of *Rhinella horribilis* in landscape 2 (P2O) as estimated with different methods. (**a**) Sampling sites in Oaxaca, southern Mexico, (**b**) SNMF (*K*=2), (**c**) TESS (*K*=2), (**d**) DAPC, (**e**) sPCA.

### Landscape genetics

Mantel test results indicated significant isolation by distance in the two landscapes (*R^2^* =0.61, *p=*0.027, P1O; *R^2^* =0.54, *p=*0.039, P2O) (S6a,b Fig). Regarding the testing of roads and rivers as barriers to gene flow, the db-RDA showed a positive relationship, supporting that a river that crosses in the north of P1O (River 2) effectively separates the north genetic group from the rest of localities (Table 2). For P2O, the MRDM showed a negative relationship in the roads model 4 (the one that considers all the roads in the landscape; S2j Fig) (*F* = -0.8236, *p=*0.015), suggesting that these roads do not limit connectivity among sampling sites. No other variables in the two landscapes were identified as relevant in explaining genetic structure (*p* >0.05).

**Table 2.**
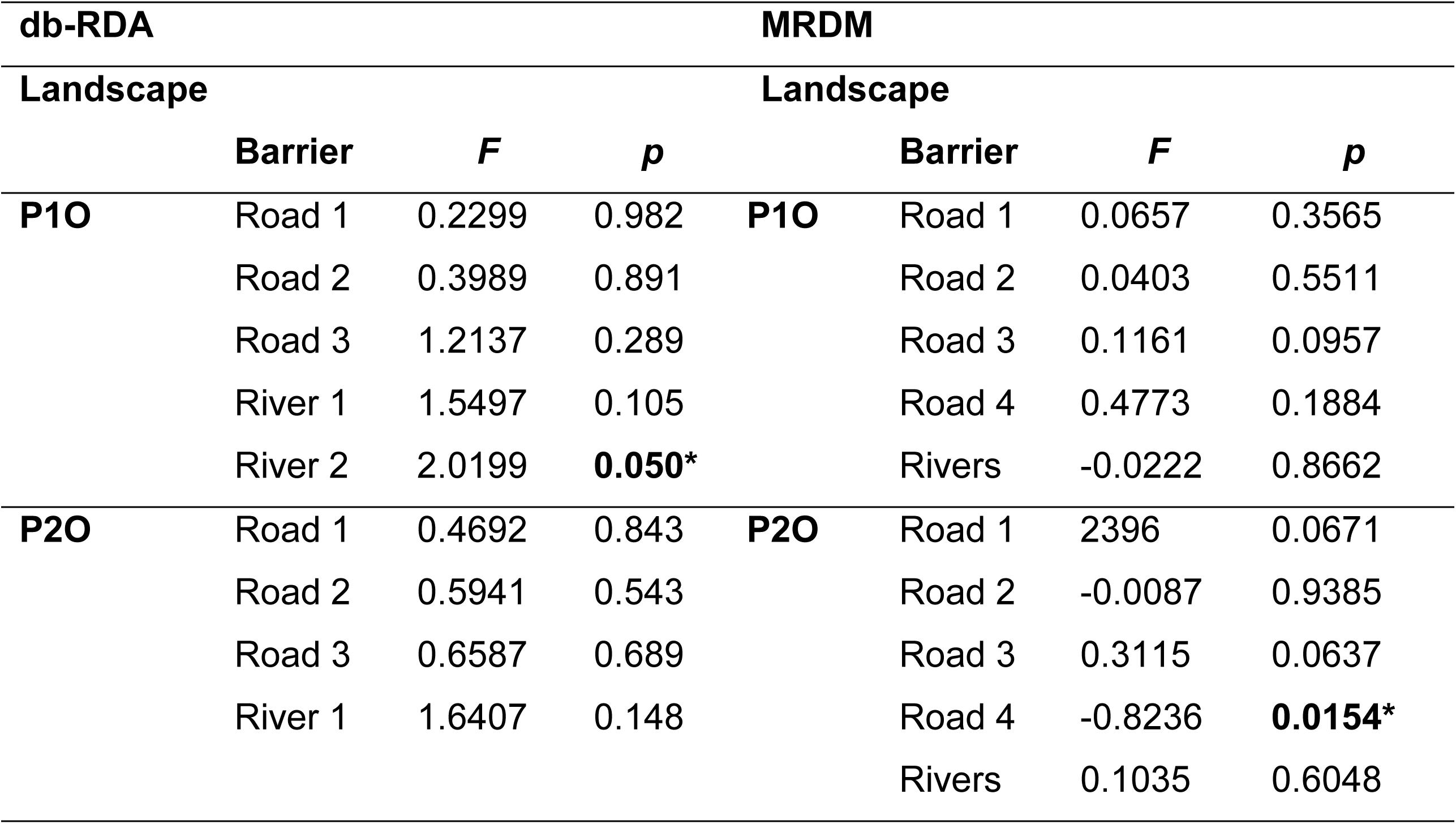
Results of isolation by barrier (IBB) models for *Rhinella horribilis* in two landscapes (P1O and P2O) in Oaxaca, southern Mexico, using distance-based RDA (db-RDA) and regression on distance matrices (MRDM). Significant results are highlighted in bold (*p* ≤ 0.05). See S2 Fig for details on each tested model.

| db-RDA |  |  |  | MRDM |  |  |  |
| --- | --- | --- | --- | --- | --- | --- | --- |
| Landscape |  |  |  | Landscape |  |  |  |
|  | Barrier | *F* | *p* |  | Barrier | *F* | *p* |
| **P1O** | Road 1 | 0.2299 | 0.982 | **P1O** | Road 1 | 0.0657 | 0.3565 |
|  | Road 2 | 0.3989 | 0.891 |  | Road 2 | 0.0403 | 0.5511 |
|  | Road 3 | 1.2137 | 0.289 |  | Road 3 | 0.1161 | 0.0957 |
|  | River 1 | 1.5497 | 0.105 |  | Road 4 | 0.4773 | 0.1884 |
|  | River 2 | 2.0199 | **0.050\*** |  | Rivers | -0.0222 | 0.8662 |
| **P2O** | Road 1 | 0.4692 | 0.843 | **P2O** | Road 1 | 2396 | 0.0671 |
|  | Road 2 | 0.5941 | 0.543 |  | Road 2 | -0.0087 | 0.9385 |
|  | Road 3 | 0.6587 | 0.689 |  | Road 3 | 0.3115 | 0.0637 |
|  | River 1 | 1.6407 | 0.148 |  | Road 4 | -0.8236 | **0.0154\*** |
|  |  |  |  |  | Rivers | 0.1035 | 0.6048 |

Model selection after optimization of univariate models showed that the temporary water bodies (TWB) was the best-supported model (85.6%; ΔAIC=0, *R^2^*=80.86) in P1O (Table 1); this model exhibited an Inverse monomolecular Ricker function (S7a,b Fig), and assigned low resistance (high connectivity) from low to intermediate probability of presence of water bodies (<0.4), with a fast increase in resistance at higher values (>0.5). The other three models had moderate support for temporary streams (TS, 7.2%; assigning low resistance values as in TWB), solar radiation (SR, 3.8%, near 18700 kJ m-2 day-1), and ambient temperature (AT, 3.4%, between 26-27°C) (S7c,d Fig). For P2O, distance (D) was the best-supported model (75.9%; ΔAIC=0, *R^2^*=61.9), followed by AT (13.48%; ΔAIC=-2.19, *R^2^*=58.13) and SR (13.48%; ΔAIC=- 2.19, *R^2^*=58.13) (Table 1), exhibiting an Inverse monomolecular and a Ricker functional forms, respectively. The AT model assigned low resistance to temperature between 20-24°C and SR values below 18600 kJ m-2 day-1 (S8a,b Fig). Relative humidity (RH; 1.7%) and TWB (1.6%) showed moderate support, assigning low resistance to high RH (>88%) and intermediate TWB (0.5-0.8) values (S8c,d Fig).

The multivariate model selection (Table 3) showed that TBW was the most-supported model (85.6%) for P1O and explained the highest variation (*R^2^*=80.91), followed by the Aquatic model (Temporary water bodies + Temporary streams; *R^2^*=7.01). In accordance, the resistance surface (Fig 4) of the Aquatic model indicated high resistance (limited connectivity) between the northernmost sampling sites (AMA-AMB) and the rest (red circle, Fig 4b), and between the central AMA-AMB (green circle) sites. Regarding P2O, in addition to distance, the most-supported models were AT (25.9%; *R^2^*=61.5), TBW (22.29%; *R^2^*=77.62) and SR (14.8%; *R^2^*=57.9) (Table 3). These results also agree with the overall patterns observed from the resistance surfaces, where TBW indicated high resistance between the genetic groups on the north, center and south (red, green, purple circles, Fig 4c), while AT and SR showed moderate resistance (Fig 4d,e).

**Fig. 4.**
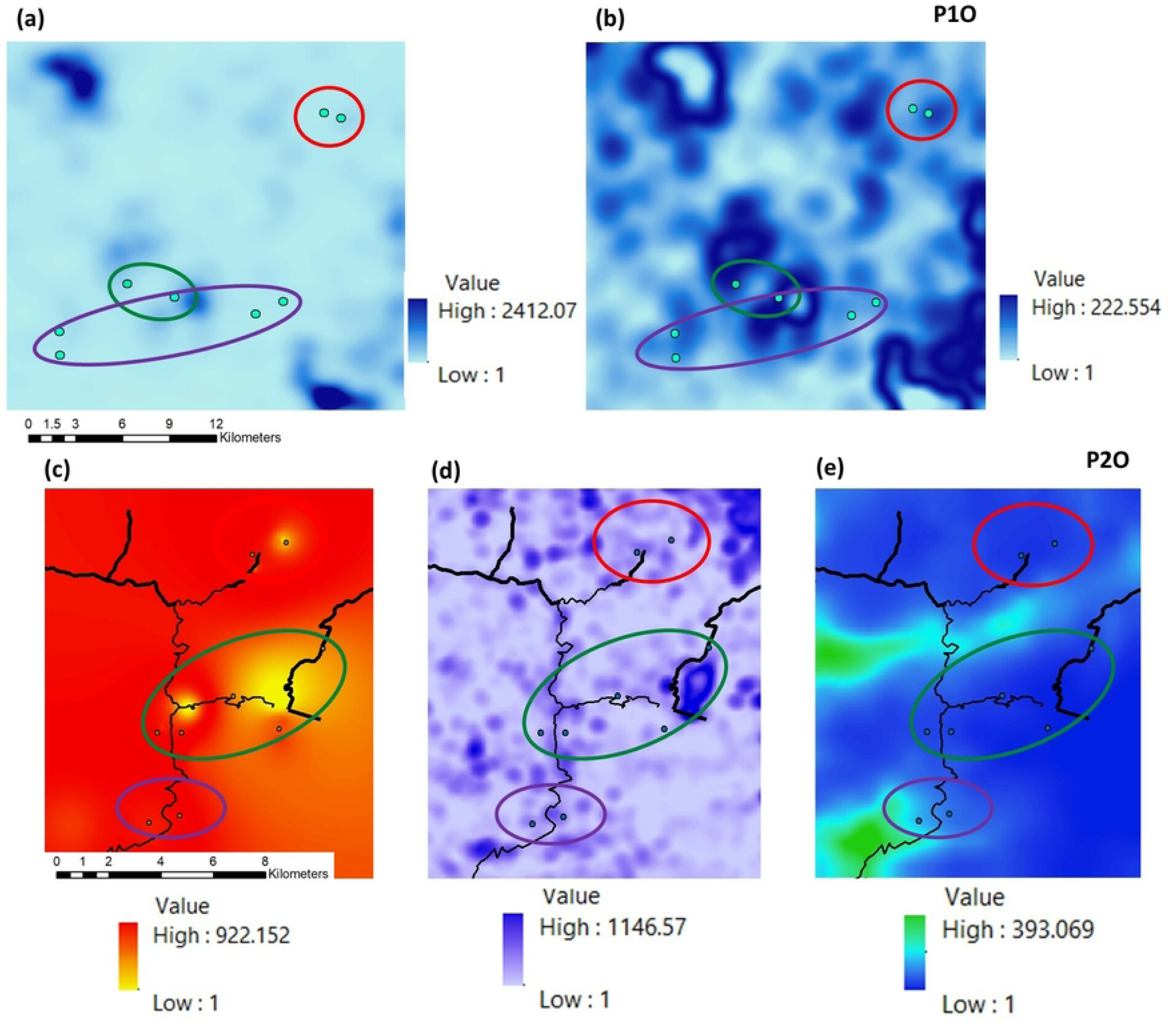
Most significant optimized resistance surfaces for *Rhinella horribilis* in two landscapes (P1O and P2O) in Oaxaca, southern Mexico (see Table 4 for details). (**a**) Temporary water bodies (TWB) and (**b**) Aquatic model (Temporary water bodies + Temporary streams) for P1O. (**c**) Ambient temperature (AT), (**d**) Aquatic model and (**e**) solar radiation (SR) for P2O. Color scales in each inset depict resistance values. Dots indicate sampling sites, and red, green and purple ovals depict the identified genetic groups.

**Table 3.**
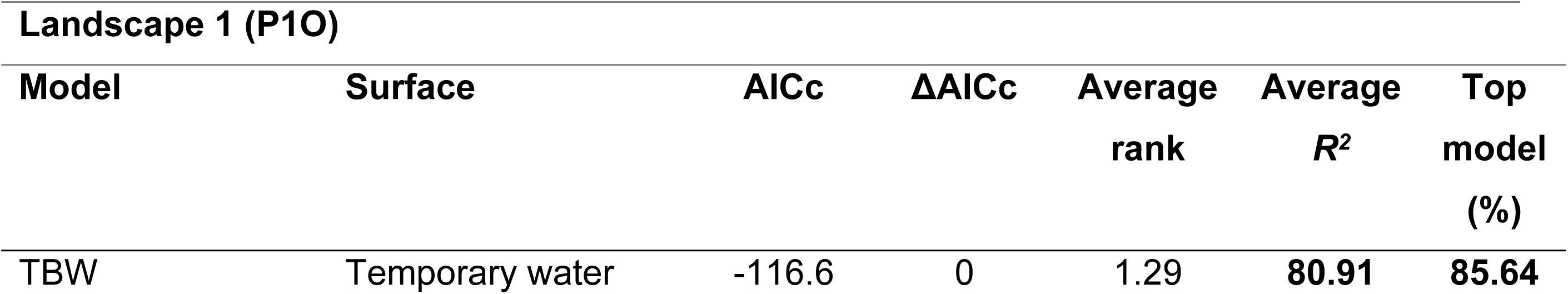
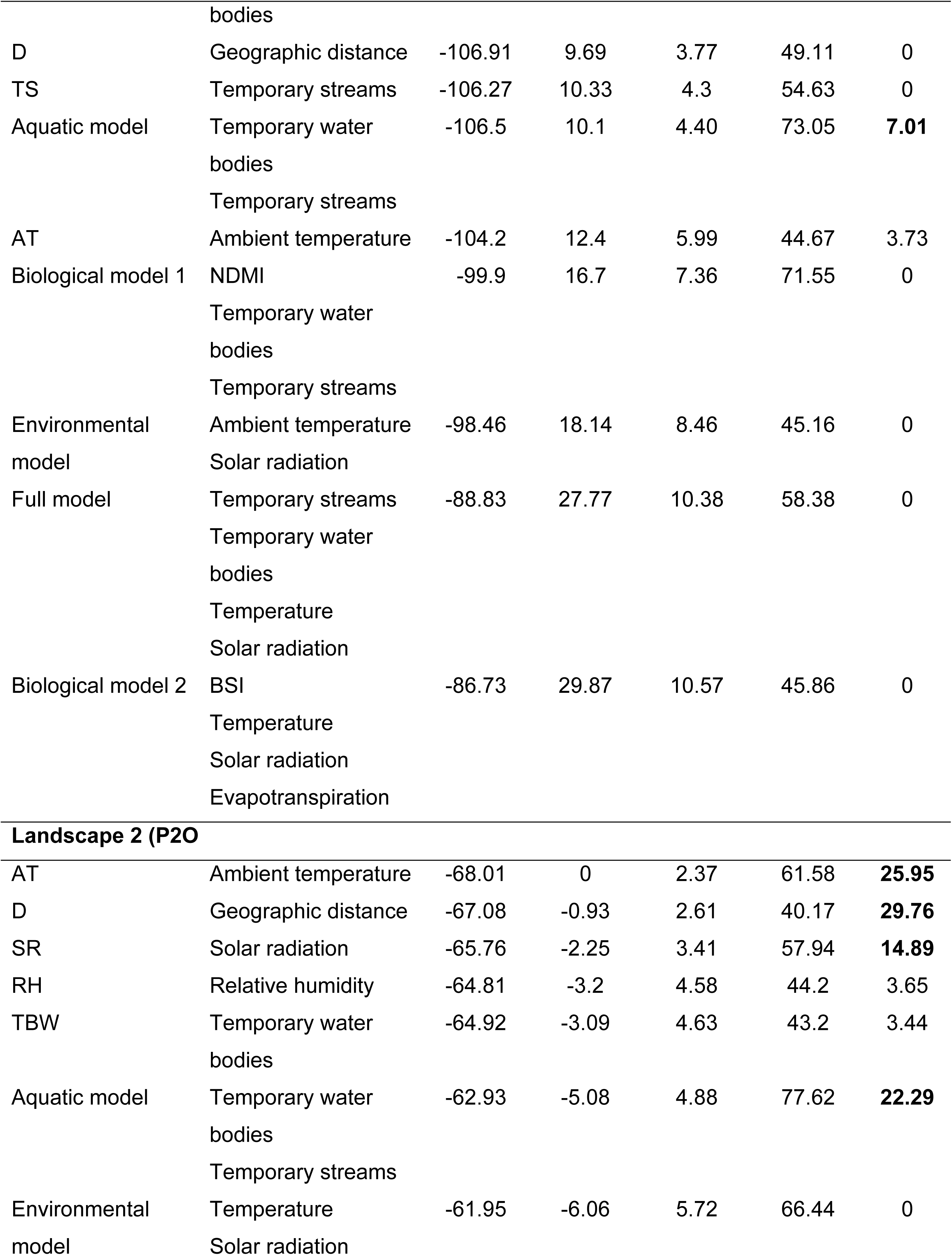
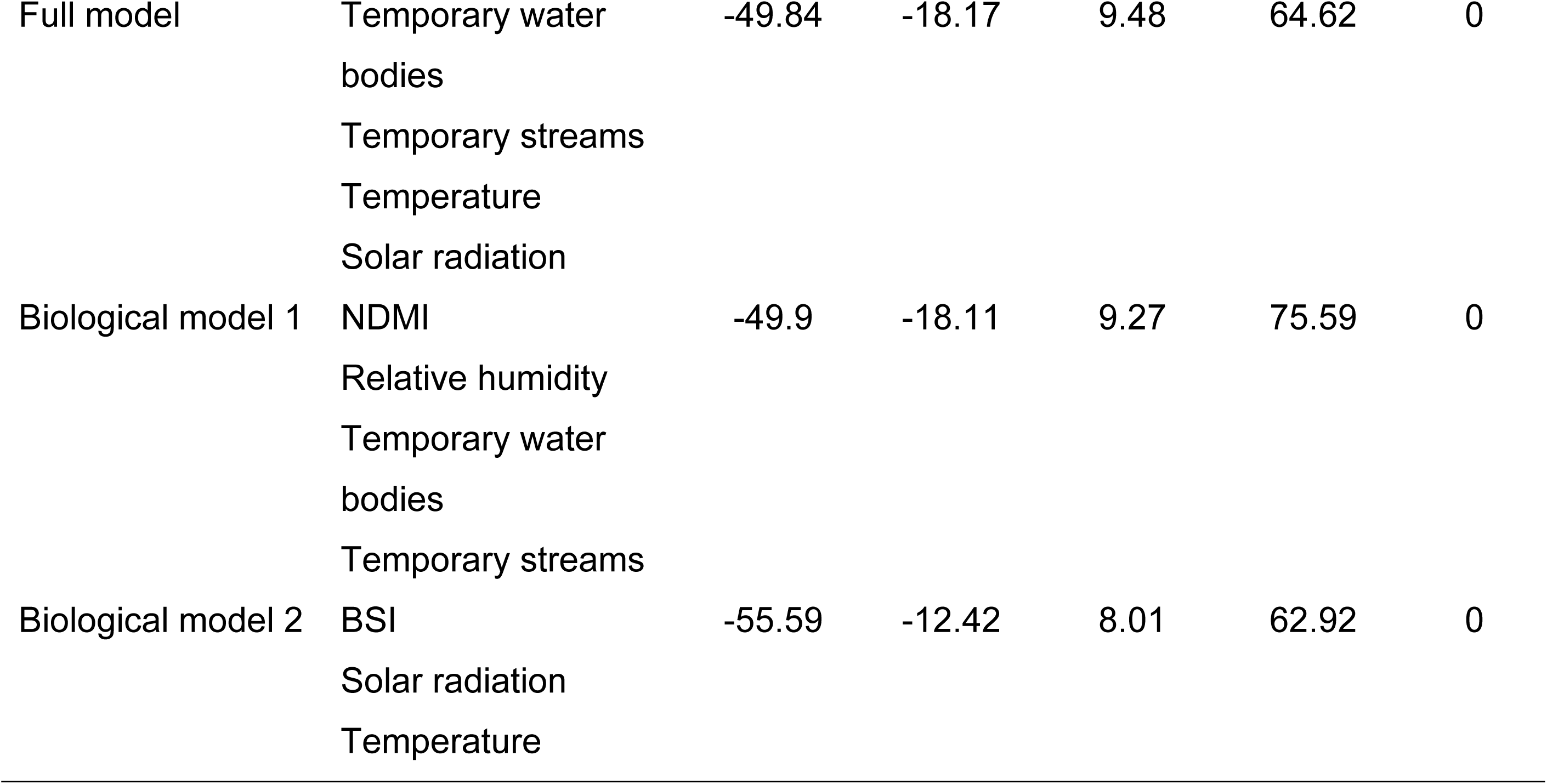
Multivariate generalized linear mixed models for *Rhinella horribilis* in two landscapes (P1O and P2O) in Oaxaca, southern Mexico. Best-supported models are indicated by highest ‘top model (%)’ in bold; AICc: Akaike information criterion. The average rank and average *R^2^* (value of the fitted model; %) are shown.

| Landscape 1 (P1O) |  |  |  |  |  |  |
| --- | --- | --- | --- | --- | --- | --- |
| Model | Surface | AICc | ΔAICc | Average rank | Average $R^2$ | Top model (%) |
| TBW | Temporary water bodies | -116.6 | 0 | 1.29 | **80.91** | **85.64** |
| D | Geographic distance | -106.91 | 9.69 | 3.77 | 49.11 | 0 |
| TS | Temporary streams | -106.27 | 10.33 | 4.3 | 54.63 | 0 |
| Aquatic model | Temporary water | -106.5 | 10.1 | 4.40 | 73.05 | **7.01** |
|  | bodies |  |  |  |  |  |
|  | Temporary streams |  |  |  |  |  |
| AT | Ambient temperature | -104.2 | 12.4 | 5.99 | 44.67 | 3.73 |
| Biological model 1 | NDMI | -99.9 | 16.7 | 7.36 | 71.55 | 0 |
|  | Temporary water |  |  |  |  |  |
|  | bodies |  |  |  |  |  |
|  | Temporary streams |  |  |  |  |  |
| Environmental model | Ambient temperature | -98.46 | 18.14 | 8.46 | 45.16 | 0 |
|  | Solar radiation |  |  |  |  |  |
| Full model | Temporary streams | -88.83 | 27.77 | 10.38 | 58.38 | 0 |
|  | Temporary water |  |  |  |  |  |
|  | bodies |  |  |  |  |  |
|  | Temperature |  |  |  |  |  |
|  | Solar radiation |  |  |  |  |  |
| Biological model 2 | BSI | -86.73 | 29.87 | 10.57 | 45.86 | 0 |
|  | Temperature |  |  |  |  |  |
|  | Solar radiation |  |  |  |  |  |
|  | Evapotranspiration |  |  |  |  |  |
| Landscape 2 (P2O) |  |  |  |  |  |  |
| AT | Ambient temperature | -68.01 | 0 | 2.37 | 61.58 | **25.95** |
| D | Geographic distance | -67.08 | -0.93 | 2.61 | 40.17 | **29.76** |
| SR | Solar radiation | -65.76 | -2.25 | 3.41 | 57.94 | **14.89** |
| RH | Relative humidity | -64.81 | -3.2 | 4.58 | 44.2 | 3.65 |
| TBW | Temporary water | -64.92 | -3.09 | 4.63 | 43.2 | 3.44 |
|  | bodies |  |  |  |  |  |
| Aquatic model | Temporary water | -62.93 | -5.08 | 4.88 | 77.62 | **22.29** |
|  | bodies |  |  |  |  |  |
|  | Temporary streams |  |  |  |  |  |
| Environmental model | Temperature | -61.95 | -6.06 | 5.72 | 66.44 | 0 |
|  | Solar radiation |  |  |  |  |  |
| Full model | Temporary water bodies | -49.84 | -18.17 | 9.48 | 64.62 | 0 |
|  | Temporary streams |  |  |  |  |  |
|  | Temperature |  |  |  |  |  |
|  | Solar radiation |  |  |  |  |  |
| Biological model 1 | NDMI | -49.9 | -18.11 | 9.27 | 75.59 | 0 |
|  | Relative humidity |  |  |  |  |  |
|  | Temporary water bodies |  |  |  |  |  |
|  | Temporary streams |  |  |  |  |  |
| Biological model 2 | BSI | -55.59 | -12.42 | 8.01 | 62.92 | 0 |
|  | Solar radiation |  |  |  |  |  |
|  | Temperature |  |  |  |  |  |

**Table 4.**
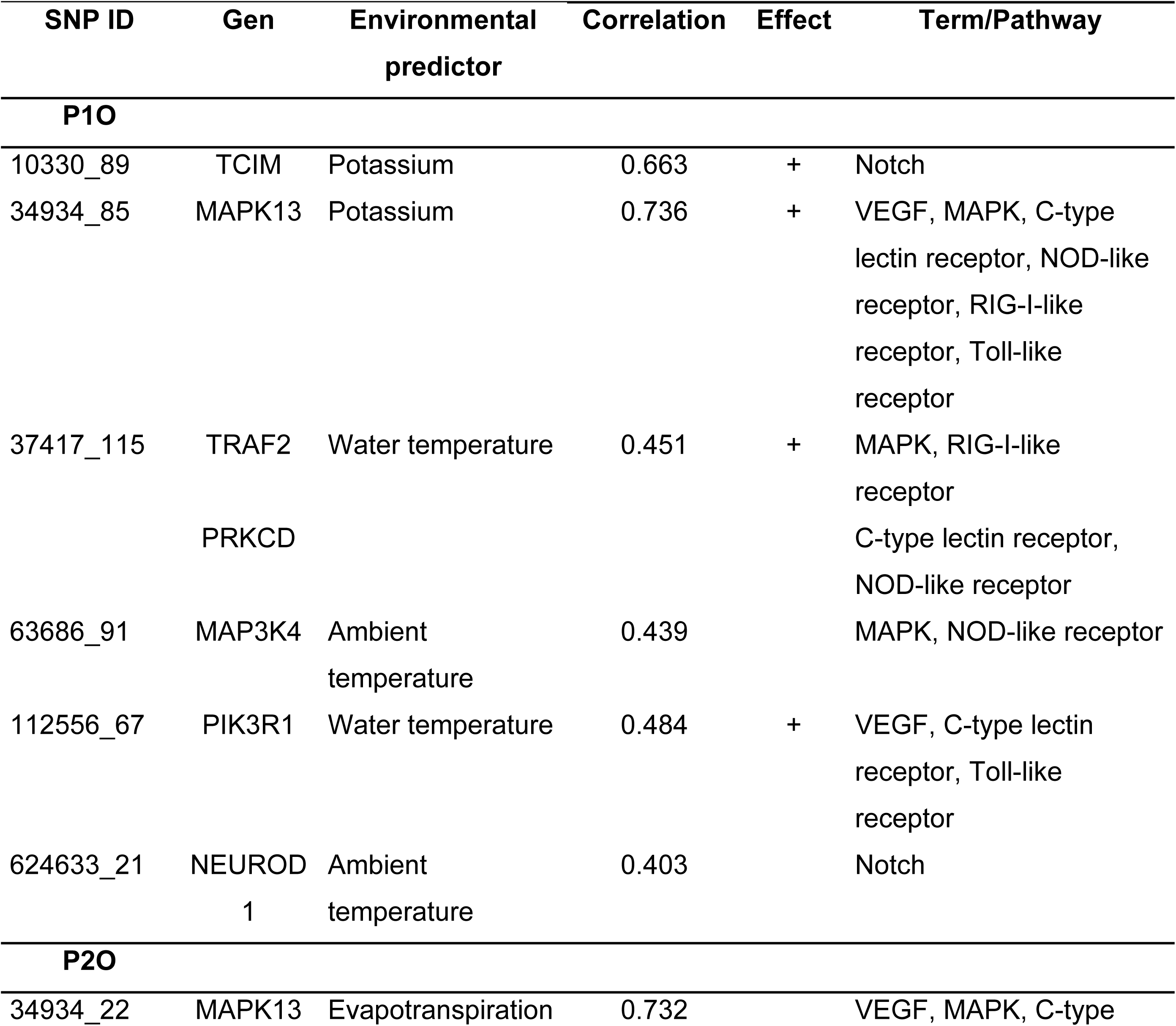
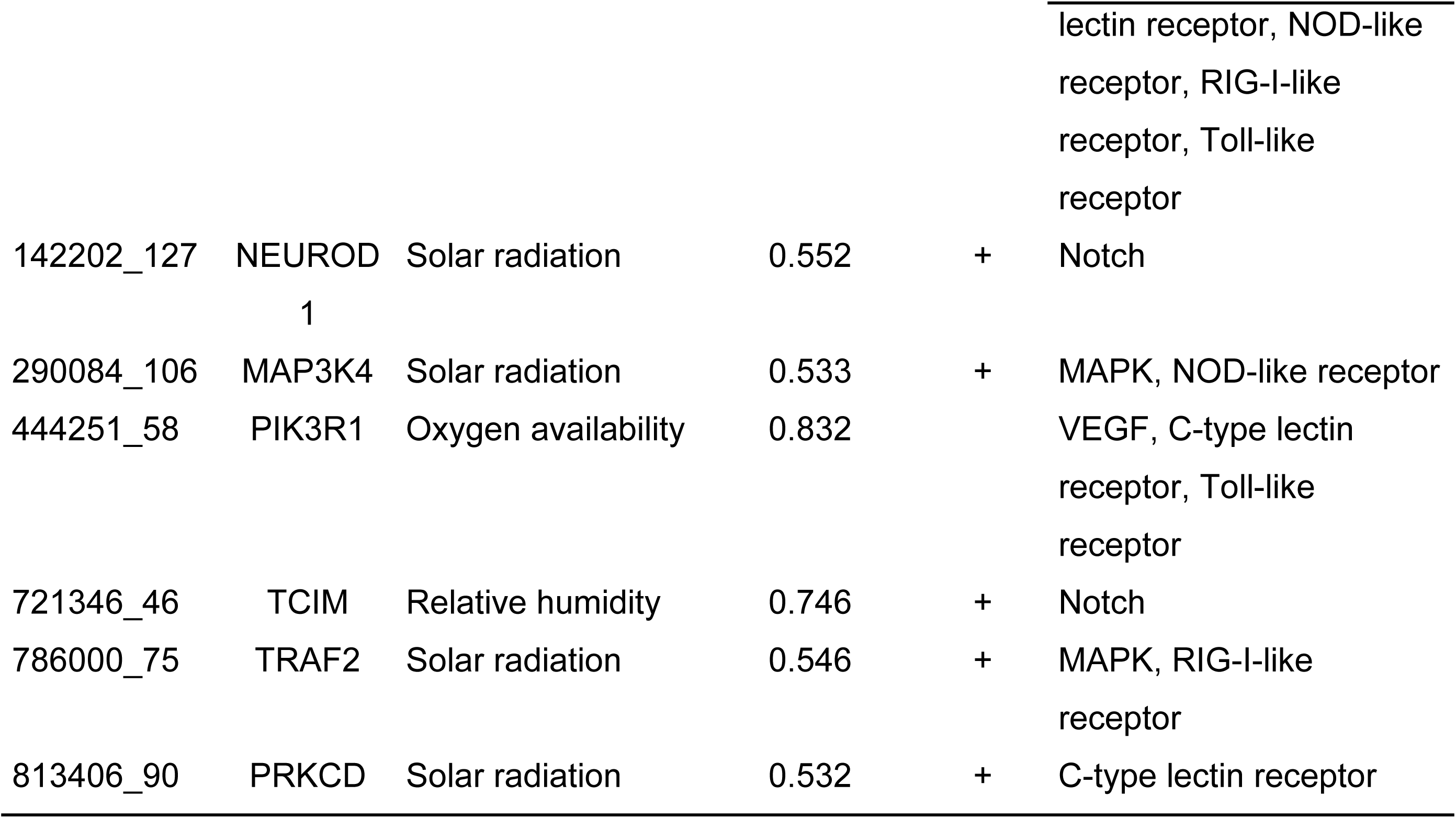
Genotype-environment associations (GEA) and corresponding GO term/KEGG pathways for *Rhinella horribilis* in two landscapes (P1O and P2O) in Oaxaca, southern Mexico. Positive (+) and negative (-) relationships are indicated.

### Identification and functional annotation of candidate loci

Results of outliers detection (candidate loci) per landscape varied by the method used; PCAdapt identified 109 and 226 unique loci in P1O and P2O, respectively; RDA 608 and 212 loci, and LFMM 2389 and 2301 loci (Table S6). Regarding environmental and physicochemical variables, LFMM identified associations with the eight variables analyzed in each landscape (Table S6); evapotranspiration (EVA) and water temperature (WT) were the variables with the highest number of associated SNPs in P1O, while in P2O these were solar radiation (SR) and oxygen availability (OA). In turn, RDA identified in P1O SNPs positively associated with potassium (K) and WT, while negatively with EVA. Associations identified in P2O were all positive, including relative humidity (RH), SR, OA and WT (Table S6). A total of 14 outlier SNPs in P1O and 2 in P2O (S9 Fig) were detected across the three methods. For cross-validating selection and reducing detection of spurious signals, we considered candidate loci the SNPs detected by at least two methods, resulting in 609 candidate SNPs for P1O and 237 for P2O, which were used for annotation and subsequent analyses. Additionally, these SNPs were homogeneously distributed across the genome (SNPs density curves and SNPs frequency; S10 Fig). We tested if there was overlap of the density curves of candidate SNPs between the two landscapes, which was not significant (*D*=0.55, test of equal densities; *p* >0.05).

The multivariate PCA results showed no differences regarding the *in situ* climatic and physicochemical variables between the two landscapes (S11 Fig). The number of candidate SNPs associated with the same environmental variables in both landscapes was significant (Fisher-test, *p <*0.05; Table S7), signaling that *R. horribilis* is likely responding similarly (not randomly) to the environmental selection pressures in the two landscapes. On the other hand, results indicated that unique processes are also occurring in P1O, since none of the 14 SNPs was associated with the same variables in the two landscapes (i.e., there is no overlap).

Furthermore, these results were supported by the RDAs performed for the different sets of SNPs, which confirm that the identified candidate SNPs best explain the landscape’s environmental variation (see Table S3 for detailed results). The candidate SNPs per landscape (Set-609-P1O and Set-237-P2O) explained between 58% and 85% of the environmental variation of their corresponding landscape, respectively (*p*=0.001), whereby the crossed tests (Set-609-P2O, Set-237-P1O) only accounted for 35% and 51% of the variation with the other landscape’s SNPs. When using the 14 SNPs detected with all three methods for P1O (Set-14- P1O), we were able to explain 61% of its environmental variation, while their association with P2O (Set-14-P2O) was lower (50%). When using both the neutral (non-correlated genetic variants from the ‘full data’) and the random SNPs sets, we could explain very low percentages of the environmental variance of both landscapes, indicating that associations were not spurious.

Functional annotation was possible for 385 regions containing the candidates detected in P1O and for 248 in P2O (E-val <0.05; Table S8), 34 of which were common to both landscapes. These 34 candidates were assigned to three main GO terms (GO:0045746, GO:0008593, GO:0005829) and 13 KEGG metabolic pathways (Table S9, Fig 5). The biological process with highest significance was the NOTCH signal (Table S9), with two SNPs in each landscape assigned to enriched genes in this signaling system (NEUROD1 and TCIM); these were associated positively with potassium (K), solar radiation (SR) and relative humidity (RH), and negatively with ambient temperature (AT) (Table 4; see Table S10 for full version). Six SNPs were assigned to cytosol cellular component. Finally, the most important KEGG metabolic pathways were the hypothalamic gonadotropin-releasing hormone (GnRH), the C-type lectin receptors (CLR), the NOD-like receptor signaling pathway, and the VEGF and MAPK signaling pathways (Table S9, Fig 5). Four and five SNPs for P1O and P2O, respectively were associated with enriched genes of these metabolic pathways, positively with SR, oxygen availability (OA), water temperature (WT) and K, and negatively with AT and evapotranspiration (Table 4; see Table S10 for full version).

**Fig. 5.**
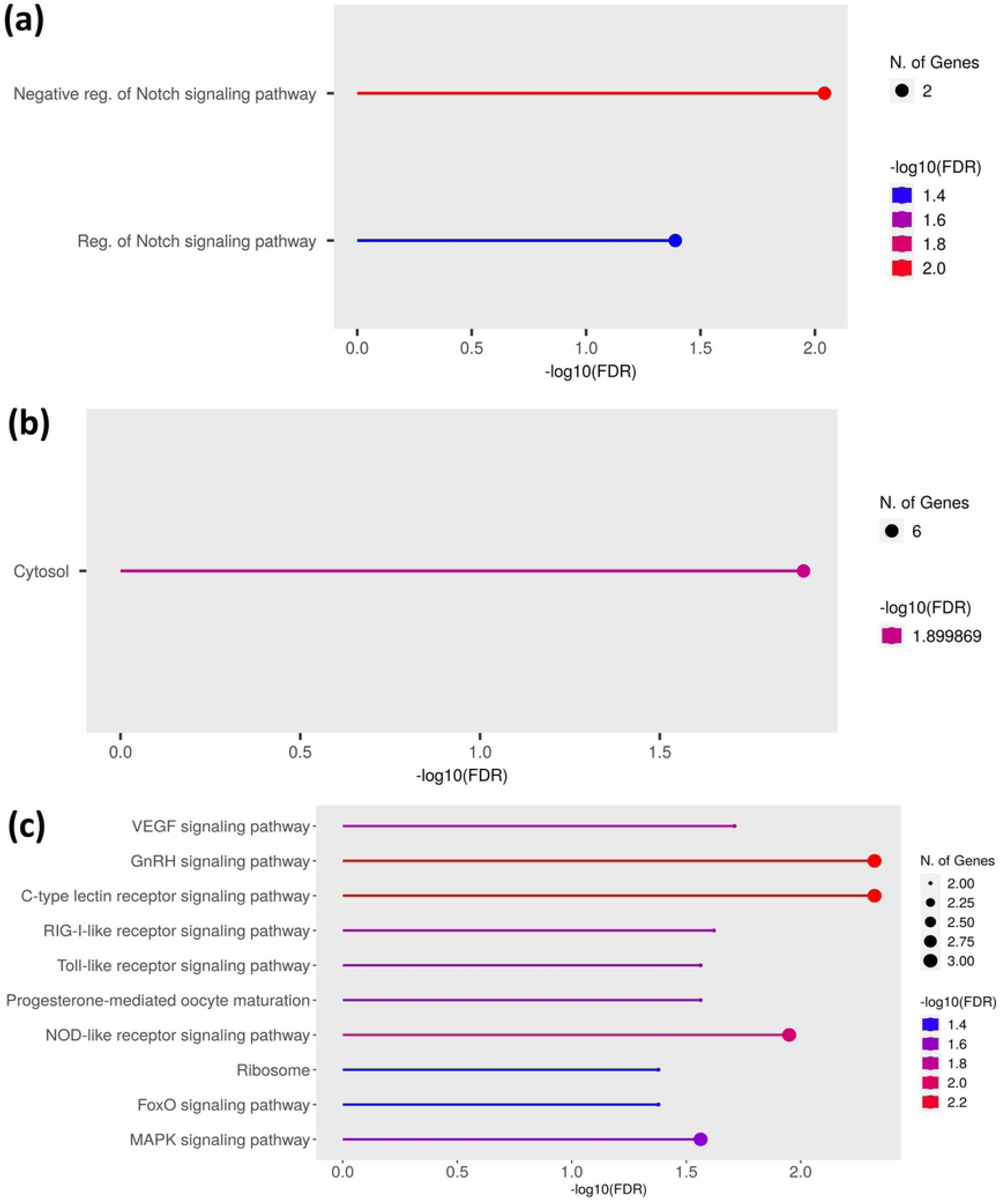
Gene enrichment analysis of the 34 SNPs common to both landscapes (P1O, P2O). Results are shown for (**a**) GO-biological process, (**b**) GO-cellular components, and (**c**) KEGG metabolic pathways, depicting the fold enrichment from highest (top) to lowest (bottom). In the case of (**c**), the selected 10 enrichment pathways with highest -log10(FDR) are included.

Enrichment analyses to identify distinct processes in each landscape, based on their unique genes, assigned more than double GO terms and KEGG metabolic pathways in P1O (228) compared with P2O (107) (FDR, *p* <0.05; Table S11, S12 Fig). Considering only the first 10 GO terms per landscape, the most important biological processes in P1O were those related to development, from early states of formation of anatomical structures to maturation, as well as genetic expression, cellular communication and cellular function (S12a Fig, Table S11). Some of the enriched molecular functions included energy transfer, transduction signals and enzymatic regulation (S12c Fig, Table S11), and three KEGG metabolic pathways were most significant, the MAPL signal, focal adhesion and MRNA surveillance pathway (S12d Fig). Comparatively, in P2O the most important biological processes and molecular functions were genetic regulation, transcription regulation and RNA biosynthesis and metabolism (S12e Fig).

## Discussion

Unlike its close relative, the amply known Cane Toad *Rhinella marina,* the Giant Toad *R. horribilis* has been little studied, particularly concerning population genetic assessments within its native distribution. Our study shows high functional connectivity associated with temporary water bodies availability, low vegetation cover, and high humidity, as well as potential signals of local adaptation to modified habitats in this species. We found both shared and unique patterns regarding landscape factors and genotype-environment associations related with the degree of modification across landscapes. Putatively adaptive SNPs were associated with genes and metabolic pathways involved in embryonic development, sexual maturation and immune responses, which may facilitate responses to the stressful habitat conditions and selective pressures imposed by anthropized habitats.

### Interplay of landscape features on genomic patterns and functional connectivity

Low levels of genetic variability and increased inbreeding are often observed in wild animal populations inhabiting fragmented and anthropogenically modified habitats [10,81], and amphibians are no exception [19,82–84]. However, we observed higher genetic diversity in the *R. horribilis* populations studied. Arruda and collaborators [85] report similar results in populations of the Cururu Toad *Rhinella schneideri* inhabiting agricultural areas; these results may be related with certain life history traits of the two species, including large body size and high dispersal ability [86], which allow them to occupy a wide variety of habitats [32].

As predicted, *R. horribilis* also showed moderate genetic structure within landscapes (P1O, P2O), which mostly displayed patterns of isolation by distance (concordant with the spatial distribution) and isolation by barrier (i.e., the main river in P1O). Amphibian species with low vagility usually exhibit more structured populations in modified compared to natural, well-preserved areas [16,17,19,84]. In contrast, species with greater mobility may counter these effects, for example cases like *R. schneideri* and the Southern Leopard Frog *Rana [Lithobates] sphenocephalus*, which show low genetic differentiation in agriculture and urban landscapes, respectively [85,87]. Interestingly, although *Rhinella* species are characterized by high dispersal capacity (up to 2 km home ranges) [88], our results show some degree of fine-scale genetic structure in *R. horribilis,* suggesting there are other factors in place influencing individual dispersal patterns.

Indeed, water bodies were the most important drivers of functional connectivity for *R. horribilis* in both landscapes. The second order river (i.e., large, strong current) that divides the northern (Santa María Zacatepec) from the rest of sampling sites in P1O limited (but not hampered) dispersal and gene flow, influencing the observed genetic structure. These results are consistent with previous studies in other amphibian species across a wide range of landscape types, showing that the magnitude, direction and intensity of water flow can have a strong impact on patterns of gene flow at the landscape scale [18,89–92].

Temporary water bodies (streams and ponds) were crucial for maintaining functional connectivity associated with the modified environment in both landscapes. Our findings showed an overall gradient where resistance was lower when water bodies were not very abundant, with resistance rapidly increasing at higher values. Low to medium density of temporary water bodies (natural and artificial), which are frequent in modified landscapes, facilitate stepping-stone dispersal. Individual dispersal among these water bodies in search of favorable conditions for shelter, feeding and reproduction would likely be reached at certain density of water bodies, facilitating genetic interchange, and likely sustaining the genetic diversity we observe in *R. horribilis* populations. These results agree with previous studies showing that temporary streams associated with riparian vegetation and forested areas promote connectivity in urban and agricultural areas (e.g., [19,92–96]). Our findings highlight the importance of temporary water bodies (ponds and streams) for dispersal and reproduction in *R. horribilis* in both landscapes, irrespective of the degree of habitat modification. Therefore, it is crucial to conserve these landscape features to maintain their population dynamics. Building artificial water bodies surrounded by vegetation cover in sites with extensive agriculture and livestock areas can be fundamental to mitigate the negative impacts of land use and climate changes on amphibian populations.

We expected that connectivity would be positively associated with low vegetation cover, secondary forest cover and high humidity, while overall high temperature and solar radiation in such areas would not limit connectivity. This was supported by our model selection results yet, interestingly, with different patterns in each landscape associated with their degree of modification. Landscape P2O is characterized by more vegetation cover, lower fragmentation and less agriculture, pastures and livestock land use areas (Fig 1). Accordingly, *R. horribilis* in P2O exhibited higher connectivity associated with moderate vegetation cover (NDVI >0.3), as well as with high relative humidity (>88%), and lower ambient temperature (AT, 20-24°C) and solar radiation (SR, <18600 kJ m-2 day-1) values; that is, the toads seem to avoid open areas with high temperature and UV exposure. Similar connectivity patterns have been documented for amphibians that occupy sites with moderate vegetation cover, including the semiaquatic salamanders *Hynobius yangi* and the terrestrial frogs *Ascaphus montanus* and *A. truei* [19,96,97]. In contrast, higher AT and SR values (26-27°C; near 18700 kJ m-2 day-1) and less vegetation cover (grassland/shrubs to bare soil) were significantly associated with connectivity in landscape P1O. Large terrestrial anurans like *R. horribilis* can use behavioral and physiological strategies to disperse through open areas and avoid water loss, using temporary water bodies and areas with high humidity.

### Genotype-environment associations and signals of adaptive genomic responses

Since *R. horribilis* can tolerate detrimental environmental and water conditions, we hypothesized that genotype-environment associations and potential adaptation would be related to the level of anthropic modification within each landscape. Our findings support this prediction; interestingly, with both shared and unique patterns in the two landscapes studied.

We identified biological processes and metabolic pathways associated with 34 genes shared in the two landscapes, which suggests parallel evolution processes [27]. Enrichment analyses revealed that these 34 shared genes were mostly involved with the NOTCH biological process. NOTCH involves regulation mechanisms that control multiple cellular differentiation processes during embryonic development, particularly neurons and muscular cells [98,99]. In fact, NOTCH was discovered in the frog *Xenopus*, in which it is active during early embryonic stages and in the development of the nervous system [100,101]. The shared genes were also involved with various KEGG metabolic pathways (MAPK, VEGF). MAPK plays an important role in complex cellular programs like proliferation, differentiation, development and apoptosis, which can be activated by growth and stress factors [102,103]. A decreased expression and function loss of the MAPK families MEK5-ERK5 and p38MAPK inhibits neuronal differentiation and causes myogenesis defects in both embryos and tadpoles in *Xenopus* [102]. In turn, VEGF modulates the angiogenesis and organogenesis during embryonic development [104–106]. Notably, anoxia is associated with an increase in NOTCH receptors and ligands in the *R. sylvatica* [107]; it is also known that hypoxia can upregulate VEGF, stimulating vein growth and increasing the VEGF mRNA average life [104]. Hence, some environmental conditions can regulate these signals, especially under stressful conditions. Likewise, GO terms and KEGG metabolic pathways were also associated with the unique genes of each landscape and, importantly, they were more than double in P1O, the landscape with the strongest habitat modification, compared with P2O. The signaling systems and pathways in these unique genes include all those identified for the shared ones (e.g., NOTCH, VEGF, MAPK, NOD-like; Table S11).

Although the function and mechanisms of the processes and pathways above are known, it is less clear which environmental factors activate and regulate them. In this regard, we detected some variables that may help us to understand which mechanisms are guiding local adaptation processes. Genes pik3r1 and mapk13, both associated with VEGF, were linked negatively and positively with oxygen availability and high potassium levels in water bodies, respectively (Table 4). A positive association was also found between TCIM, NEUROD1, TRAF2, and MAP3K4 with high temperature and potassium levels. Low water oxygen levels affect survival and the development of tadpoles [108], while high temperatures promote metamorphosis but at reduced size, as shown in other bufonid toads (e.g., *Epidalea calamita* [109] and *Bufo gargarizans* [110]). Likewise, high levels of potassium and sodium ions negatively affect hatching, muscle development and survival in embryos and tadpoles [108,111]. Nonetheless, *Rhinella granulosa* and *R. marina* develop adequately in similar aquatic conditions [112,113].

Small or temporary water bodies in anthropogenic landscapes commonly have high temperatures, high levels of pollutants (e.g., pesticides, sodium, potassium), hypoxic conditions and shorter hydroperiods, with negative impacts on amphibian metamorphosis [3,108,114].

*Rhinella horribilis* is an explosive breeder in the temporary water bodies studied, which formed at the beginning of the rainy season. Many of these ponds were small, had detrimental water conditions, and dried fast, hence exerting selective pressures for eclosion and metamorphosis to occur rather rapidly. In such conditions, upregulated NOTCH and MAPK could be crucial for the rapid development of embryos and tissue formation, and for maintaining stable osmotic conditions in larvae. Similarly, upregulated VEGF could facilitate gas exchange and faster organ development during metamorphosis. Indeed, Wang et al. [115] found similar enriched VEGF and MAPK pathways in the fish *Leuciscus waleckii* inhabiting alkaline and hypoxic water bodies.

While the successful reproduction of *R. horribilis* in the studied landscapes might suggest local adaptation, further studies in controlled conditions are needed to confirm these observations.

We also found a positive association between the NEUROD1 gene (present in NOTCH) and solar radiation (Table 4). Solar radiation might have a strong impact in skin homeostasis, which is also modulated by the NOTCH signaling pathway, by promoting keratinocyte differentiation and darker skin pigmentation as a response to UV radiation [116,117].

Amphibians have behavioral, physiological and molecular mechanisms for UV radiation protection and different skin colors represent adaptation strategies to survive intense UV radiation [118,119]. For instance, Franco-Belussi & Olvera [120] found an increase of skin melanocytes in the Cuyaba Dwarf Frog *Physalaemus nattereri* after exposure to UV radiation, and dark skin phenotypes in the Plateau Brown Frog *Rana kukunoris* have upregulated genes associated with biosynthesis of melanosomes [117]. Hence, the NOTCH signaling participating in tissue repair could play a protective role for the keratinocytes enriched with melanosomes [98,119,121], potentially enabling *R. horribilis* to tolerate higher solar radiation in the most anthropized landscapes like P1O.

The skin is the foremost physical barrier of amphibians against external factors, and it is where the immune system is activated via the skin’s permeability, the mucosal antibodies, the microbiome and the antimicrobial peptides [122–124]. When these defense systems are evaded, the Pattern Recognition Receptors (PRRs) are activated, which are part of the innate immune system; their function is to recognize infectious agents and to activate immune, microbicidal, pro-inflammatory and antiviral responses, and apoptosis [122,124–128]. Notably, we found enrichment of metabolic pathways associated with these receptors, including the NOD-like, the C-type lectin and Toll-like receptors that specialize in recognition of fungal beta glucanes [126,129], as well as the RIG-I-like for viral RNA recognition [128,130]. The amphibian’s immune response is directly influenced by environmental stressors, like UV radiation [119,131], and different genes in amphibians have been associated with immune responses [132] exposed to bacteria, virus and pathogens like the fungus *Batrachochytrium dendrobatidis* (Bd) [133,134], and in invasive populations like *R. marina* [135–137], among other examples.

Moreover, we found PRRs genes (TRAF2, PIK3R1, MAPK13 and PRCD) associated with water bodies that exhibited high temperatures and alkaline and anoxic conditions (Table 4).

Pathogens proliferate in water bodies with detrimental conditions, which also modify the amphibian’s microbiome, increasing the probability of infection [27,121,138–140]. Although PRRs have been little studied in amphibians, it is known that early activation and regulation of PPRs for pathogen detection is fundamental to avoid infections in embryonic, larval and adult life stages. The RIG-I gene (present in the RIG-I-like receptor pathway) is upregulated in kidney and spleen tissues infected with iridovirus in the Chinese Giant Salamander *Andrias davidianus* [141]. Toll-like receptors are associated with antiviral and antibacterial immunity in the Dybovsky’s Frog *Rana dybowskii* [127]. Upregulated genes of the Toll-like and NOD-like receptors associated with the immune system are known in American Bullfrogs (*Rana [Lithobates] catesbeianus*) infected with the bacterium *Elizabethkingia miricola* [142]. Notably, although evidence of the PRRs role in Bd response is incipient, the PRRs detection of pathogen antigens is crucial for lymphocyte recognition by the adaptative immune system [122]. Indeed, the toad *Brachycephalus pitanga* presents upregulated genes associated with T-cell receptors in early stages of Bd infection [133]. The interaction between the innate and adaptative immune systems is likely related with a rapid and effective response when exposed a second time to the pathogen; namely, the PRRs can be subject of natural selection against infections system [122]. These genes may also help *R. horribilis* occupy water bodies with suboptimal conditions and outcompete other local amphibians.

## Conclusions and future directions

Anthropized environments are challenging for wild amphibian populations, and identifying cases of local adaptation can illuminate how populations deal with those stressful conditions, as illustrated by the neutral and adaptive patterns observed in *R. horribilis*. Comparative studies on syntopic species should be conducted to assess if they respond similarly at the genomic level to analogous selection pressures, and to characterize key landscape features for functional connectivity and survival. Further, while we found signals of parallel evolution in *R. horribilis* in the two landscapes studied, representing different degrees of human modification of their natural habitats, it would be interesting to expand the study to include populations from areas with no anthropogenic impacts, to assess if the genetic variants identified are fixed or specifically associated to these environments but absent from natural (conserved) ones. Also, common garden experiments could be designed to assess the interplay of these genes and metabolic pathways and the effects of environmental variables on morphological characteristics and in the survival of tadpoles. The identified genes and metabolic pathways and their associated functions can prove crucial for amphibian adaptation to anthropized environments, allowing individuals to tolerate adverse environmental and water conditions, as suggested by our results. In this regard, it would be interesting to evaluate the expression of upregulated genes associated with these pathways and the molecular mechanisms acting on tadpole development and survival in mesocosm experiments.

## Acknowledgements

We are grateful to R. Palacios, H. Colín, S. Hernández, R. Peralta and R. López for their enthusiastic help during fieldwork, and to N. Gálvez-Reyes for providing molecular laboratory assistance. We deeply thank the people from the field sites in Oaxaca for allowing us to work on their ejidos and private properties. Gerardo J. Soria-Ortiz acknowledges that this paper was a part of his doctoral thesis in the Programa de Doctorado de Ciencias Biológicas de la Universidad Nacional Autónoma de México (UNAM) and received a scholarship provided by the Consejo Nacional de Ciencia y Tecnología (CONACyT CVU: 814369/No. Beca: 464869), and support from Programa de Estudios de Posgrado (PAEP) and UNAM. Open access was obtained thanks to the UNAM agreement that covers article publication charges.

## Supporting information

### Supporting figures S1-S12 in one pdf file

**S1. Fig.** DAPC cross-validation for number of axes retained

**S2. Fig.** Surfaces tested for main roads and rivers used in MRDM

**S3. Fig.** Redundancy analysis (RDA) testing scheme using different sets of SNPs

**S4. Fig.** Results of structure analysis in landscape 1 (P1O)

**S5. Fig.** Results of structure analysis in landscape 2 (P2O)

**S6. Fig.** Isolation by distance plots for landscapes P1O and P2O

**S7. Fig.** Resistance curves resulting from surface optimization in P1O

**S8. Fig.** Resistance curves resulting from surface optimization in P2O

**S9. Fig.** Venn diagram of the outlier loci identified for *Rhinella horribilis* with three methods (RDA, LFMM and PCAdapt)

**S10. Fig.** Density curves and frequency of the identified candidate SNPs for *Rhinella horribilis*

**S11. Fig.** PCA of the environmental variables within each landscape (P1O and P2O)

**S12. Fig.** GO terms (GO) and KEGG metabolic pathways of the unique genes of each landscape (P1O and P2O)

### Supporting tables S1-S9 in individual files

**Table S1.** Landscape variables used for connectivity/selection models for *Rhinella horribilis*

**Table S2.** Hypotheses of multivariate models tested

**Table S3.** Results of redundancy analysis tests with the different sets of SNPs

**Table S4.** Genetic diversity results for *Rhinella horribilis* in P1O and P2O

**Table S5.** Results of paired *Fst* between sampling sites in P1O and P2O

**Table S6.** Outlier loci detected for *Rhinella horribilis* with three methods (RDA, LFMM and PCAdapt)

**Table S7.** Fisher-test results of the overlap of SNPs associated with the same *in situ* environmental variables in P1O and P2O

**Table S8.** Gene enrichment results identified for 34 shared genes in P1O and P2O

**Table S9.** Results of redundancy analysis tests for the association between enriched SNPs and the landscape variables in P1O and P2O

## References

1. Hendry AP 2017. Eco-evolutionary Dynamics. Princeton: Princeton University Press; 2017. doi: 10.1515/9781400883080

2. Lynch M. The evolution of genetic networks by non-adaptive processes. Nature Reviews Genetics. 2007; 8(10):803–813. doi: 10.1038/nrg2192

3. Covarrubias S, González C, Gutiérrez-Rodríguez C. Effects of natural and anthropogenic features on functional connectivity of anurans: a review of landscape genetics studies in temperate, subtropical and tropical species. Journal of Zoology. 2021; 313(3):159–171. doi: 10.1111/jzo.12851

4. Exposito-Alonso M, Booker TR, Czech L, Gillespie L, Hateley S, Kyriazis CC, et al. Genetic diversity loss in the Anthropocene. Science. 2022; 377(6613):1431–1435. doi: 10.1126/science.abn5642.

5. Vázquez-Domínguez E, Kassen R, Schroer S, De Meester L, Johnson MT. Recentering evolution for sustainability science. Global Sustainability. 2024; 7(e8):1–6 doi: 10.1017/sus.2024.5

6. Bar-Massada A, Radeloff VC, Stewart SI. Biotic and abiotic effects of human settlements in the wildland-urban interface. Bioscience. 2014; 64(5):429–437. doi: 10.1093/biosci/biu039

7. Bounas A, Keroglidou M, Toli EA, Chousidis I, Tsaparis D, Leonardos I, Sotiropoulos K. Constrained by aliens, shifting landscape, or poor water quality? Factors affecting the persistence of amphibians in an urban pond network. Aquatic Conservation: Marine and Freshwater Ecosystems. 2020; 30(5):1037–1049. doi: 10.1002/aqc.3309

8. Lenhardt PP, Brühl CA, Leeb C, Theissinger K. Amphibian population genetics in agricultural landscapes: does viniculture drive the population structuring of the European common frog (*Rana temporaria*)? PeerJ. 2017; 5:e3520. doi: 10.7717/peerj.3520

9. Leigh DM, Hendry A, Vázquez-Domínguez E, Friesen V. Estimated six percent loss of genetic variation in wild populations since the Industrial Revolution. Evolutionary Applications. 2019; 12(8):1505–1512. doi:10.1111/eva.12810

10. Miles LS, Rivkin LR, Johnson MT, Munshi-South J, Verrelli BC. Gene flow and genetic drift in urban environments. Molecular Ecology. 2019; 28(18):4138–4151. doi: 10.1111/mec.15221

11. Crates R, Olah G, Adamski M, Aitken N, Banks S, Ingwersen D, et al. Genomic impact of severe population decline in a nomadic songbird. PLoS ONE. 2019; 14(10):e0223953. doi: 10.1371/journal.pone.0223953

12. Oziolor EM, Reid NM, Yair S, Lee KM, Guberman VerPloeg S, Bruns PC, et al. Adaptive introgression enables evolutionary rescue from extreme environmental pollution. Science. 2019; 364(6439):455–457. doi: 10.1126/science.aav4155

13. Soria-Ortiz GJ, Ochoa-Ochoa LM, Vázquez-Domínguez E. Respuesta genética y funcional de anfibios a perturbaciones causadas por actividades antrópicas. Ecosistemas. 2023; 32(1):2462–2462. doi: 10.7818/ECOS.2462

14. Ruiz-Miñano M, While GM, Yang W, Burridge CP, Salvi D, Uller T. Population genetic differentiation and genomic signatures of adaptation to climate in an abundant lizard. Heredity. 2022; 128(4):271–278. doi: 10.1038/s41437-022-00518-0

15. Tigano A, Friesen VL. Genomics of local adaptation with gene flow. Molecular Ecology. 2016; 25(10):2144–2164. doi: 10.1111/mec.13606

16. Wenner SM, Murphy MA, Delaney KS, Pauly GB, Richmond JQ, Fisher RN, Robertson JM. Natural and anthropogenic landscape factors shape functional connectivity of an ecological specialist in urban Southern California. Molecular Ecology. 2022; 31(20):5214–5230. doi: 10.1111/mec.16656

17. Unglaub B, Cayuela H, Schmidt BR, Preißler K, Glos J, Steinfartz S. Context-dependent dispersal determines relatedness and genetic structure in a patchy amphibian population. Molecular Ecology. 2021; 30(20):5009–5028. doi: 10.1111/mec.16114

18. Homola JJ, Loftin CS, Kinnison MT. Landscape genetics reveals unique and shared effects of urbanization for two sympatric pool-breeding amphibians. Ecology and Evolution. 2019; 9(20):11799–11823. doi: 10.1002/ece3.5685

19. Jeon JY, Jeong D, Borzée A, Heo K, Park HC, Lee H, Min MS. Variation in functional connectivity between metapopulations in urbanized and forested areas in an endangered salamander. Urban Ecosystems. 2024; 27:111–124. doi: 10.1007/s11252-023-01434-9

20. Fusco NA, Pehek E, Munshi-South J. Urbanization reduces gene flow but not genetic diversity of stream salamander populations in the New York City metropolitan area. Evolutionary Applications. 2021; 14(1):99–116. doi: 10.1111%2Feva.13025

21. Van Buskirk J, Jansen van Rensburg A. Relative importance of isolation-by-environment and other determinants of gene flow in an alpine amphibian. Evolution. 2020; 74(5):962–978. doi: 10.1111/evo.13955

22. Homola JJ, Loftin CS, Cammen KM, Helbing CC, Birol I, Schultz TF, Kinnison MT. Replicated landscape genomics identifies evidence of local adaptation to urbanization in wood frogs. Journal of Heredity. 2019; 110(6):707–719. doi: 10.1093/jhered/esz041

23. Osterberg JS, Cammen KM, Schultz TF, Clark BW, Di Giulio RT. Genome-wide scan reveals signatures of selection related to pollution adaptation in non-model estuarine Atlantic killifish (*Fundulus heteroclitus*). Aquatic Toxicology. 2018; 200:73–82. doi: 10.1016/j.aquatox.2018.04.017

24. Harris SE, Munshi-South J. Signatures of positive selection and local adaptation to urbanization in white-footed mice (*Peromyscus leucopus*). Molecular Ecology. 2017; 26(22):6336–6350. doi: 10.1111/mec.14369

25. Perrier C, Lozano del Campo A, Szulkin M, Demeyrier V, Gregoire A, Charmantier A. Great tits and the city: Distribution of genomic diversity and gene-environment associations along an urbanization gradient. Evolutionary Applications. 2018; 11(5):593–613. doi: 10.1111/eva.12580

26. Winchell KM, Campbell-Staton SC, Losos JB, Revell LJ, Verrelli BC, Geneva AJ. Genome- wide parallelism underlies contemporary adaptation in urban lizards. Proceedings of the National Academy of Sciences. 2023; 120(3):e2216789120. doi: 10.1073/pnas.2216789120

27. Pabijan M, Palomar G, Antunes B, Antoł W, Zieliński P, Babik W. Evolutionary principles guiding amphibian conservation. Evolutionary Applications. 2020; 13(5):857–878. doi: 10.1111/eva.12940

28. Skelly DK, Joseph LN, Possingham HP, Freidenburg LK, Farrugia TJ, Kinnison MT, Hendry AP. Evolutionary responses to climate change. Conservation Biology. 2007; 21:1353–1355. https://www.jstor.org/stable/4620959

29. Savage AE, Zamudio KR. Adaptive tolerance to a pathogenic fungus drives major histocompatibility complex evolution in natural amphibian populations. Proceedings of the Royal Society B: Biological Sciences. 2016; 283(1827):20153115. doi: 10.1098/rspb.2015.3115

30. De León LF, Castillo A. *Rhinella marina* (Cane toad). Salinity tolerance. Herpetological Review. 2015; 46:237–238.

31. Ramírez-Arce DG, Ochoa-Ochoa LM, Lira-Noriega A. Effect of landscape composition and configuration on biodiversity at multiple scales: a case study with amphibians from Sierra Madre del Sur, Oaxaca, Mexico. Landscape Ecology. 2022; 37(8):1973–1986. doi: 10.1007/s10980-022-01479-9

32. Cortés-Suárez JE. Uso de microhábitat por parte del sapo gigante *Rhinella horribilis* en pastizales en el municipio de Villa de Leyva, Boyacá, Colombia. Revista Biodiversidad Neotropical. 2017; 7(4):253–257. doi: 10.18636/bioneotropical.v7i4.701

33. Sinsch U. Movement ecology of amphibians: from individual migratory behaviour to spatially structured populations in heterogeneous landscapes. Canadian Journal of Zoology. 2014; 92(6):491–502. doi: 10.1139/cjz-2013-0028

34. Richardson JL. Divergent landscape effects on population connectivity in two co-occurring amphibian species. Molecular Ecology. 2012; 21(18):4437–4451. doi: 10.1111/j.1365-294x.2012.05708.x

35. Beaupre SJ, Jacobson ER, Lillywhite HB, Zamudio KR. Guidelines for use of live amphibians and reptiles in field and laboratory research. 2nd Edition. Revised by the Herpetological Animal Care and Use Committee (HACC) of the American Society of Ichthyologists and Herpetologists, Lawrence, Kansas; 2004.

36. Campton BW, McGarigal K, Cushman SA, Gamble LR. A resistant-kernel model of connectivity for amphibians that breed in vernal pools. Conservation Biology. 2007; 21(3):788–799. doi: 10.1111/j.1523-1739.2007.00674.x

37. QGIS, Development Team. QGIS Geographic Information System. Open Source Geospatial Foundation Project; 2021. http://qgis.osgeo.org

38. Lepais O, Weir JT. Sim RAD: an R package for simulation-based prediction of the number of loci expected in RAD seq and similar genotyping by sequencing approaches. Molecular Ecology Resources. 2014; 14(6):1314–1321. doi: 10.1111/1755-0998.12273

39. R Core Team. R: A language and environment for statistical computing. R Foundation for Statistical Computing; 2022.

40. Peterson BK, Weber JN, Kay EH, Fisher HS, Hoekstra HE. Double digest RADseq: an inexpensive method for de novo SNP discovery and genotyping in model and non-model species. PloS ONE. 2012; 7(5):e37135. doi: 10.1371/journal.pone.0037135

41. Catchen J, Hohenlohe PA, Bassham S, Amores A, Cresko WA. Stacks: an analysis tool set for population genomics. Molecular Ecology. 2013; 22(11):3124–3140. doi: 10.1111/mec.12354

42. Andrews S. FastQC: a quality control tool for high throughput sequence data; 2010. https://www.bioinformatics.babraham.ac.uk/projects/fastqc

43. Bolger AM, Lohse M, Usadel B. Trimmomatic: a flexible trimmer for Illumina sequence data. Bioinformatics. 2014; 30(15):2114–2120. doi: 10.1093/bioinformatics/btu170

44. Li H, Durbin R. Fast and accurate long-read alignment with Burrows-Wheeler transform. Bioinformatics. 2010; 26:589–595.

45. Danecek P, Auton A, Abecasis G, Albers CA, Banks E, DePristo MA, et al. 1000 Genomes Project Analysis Group. The variant call format and VCFtools. Bioinformatics. 2011; 27(15):2156–2158. https://www.ncbi.nlm.nih.gov/pmc/articles/PMC3137218/

46. Aguirre-Liguori JA, Ramírez-Barahona S, Gaut BS. The evolutionary genomics of species’ responses to climate change. Nature Ecology and Evolution. 2021; 5:1350–1360. doi: 10.1038/s41559-021-01526-9

47. Benestan LM, Ferchaud AL, Hohenlohe PA, Garner BA, Naylor GJP, Baums IB, et al. Conservation genomics of natural and managed populations: building a conceptual and practical framework. Molecular Ecology. 2016; 25:2967–2977. doi: 10.1111/mec.13647

48. Jombart T. adegenet: a R package for the multivariate analysis of genetic markers. Bioinformatics. 2008; 24(11):1403–1405. doi: 10.1093/bioinformatics/btn129

49. Kamvar ZN, Tabima JF, Grünwald NJ. Poppr: an R package for genetic analysis of populations with clonal, partially clonal, and/or sexual reproduction. PeerJ. 2014; 2:e281 doi: 10.7717/peerj.281

50. Latorre-Cárdenas MC, Gutiérrez-Rodríguez C, Rico Y, Martínez-Meyer E. Do landscape and riverscape shape genetic patterns of the Neotropical otter, *Lontra longicaudis*, in eastern Mexico? Landscape Ecology. 2021; 36:69–87. doi: 10.1007/s10980-020-01114-5

51. Frichot E, Mathieu F, Trouillon T, Bouchard G, François O. Fast and efficient estimation of individual ancestry coefficients. Genetics. 2014; 196(4):973–983. doi: 10.1534/genetics.113.160572

52. Caye K, Deist TM, Martins H, Michel O, François O. TESS3: fast inference of spatial population structure and genome scans for selection. Molecular Ecology Resources. 2016; 16(2):540–548. doi: 10.1111/1755-0998.12471

53. Goudet J. Hierfstat, a package for R to compute and test hierarchical F-statistics. Molecular Ecology Notes. 2005; 5(1):184–186. doi: 10.1111/j.1471-8286.2004.00828.x

54. Vavrek MJ, Vavrek MMJ. Package ‘fossil’. Palaeoecological and palaeogeographical analysis tools; 2012. https://palaeo-electronica.org/2011_1/238/index.html

55. Rogic A, Tessier N, Legendre P, Lapointe FJ, Millien V. Genetic structure of the white-footed mouse in the context of the emergence of Lyme disease in southern Quebec. Ecology and Evolution. 2013; 3(7):2075–2088. doi: 10.1002/ece3.620

56. Legendre P, Anderson MJ. Distance-based redundancy analysis: testing multispecies responses in multifactorial ecological experiments. Ecological Monographs. 1999; 69(1):1–24. doi: 10.1890/0012-9615(1999)069[0001:DBRATM]2.0.CO;2

57. Kierepka EM, Anderson SJ, Swihart RK, Rhodes Jr OE. Evaluating the influence of life-history characteristics on genetic structure: a comparison of small mammals inhabiting complex agricultural landscapes. Ecology and Evolution. 2016; 6(17):6376–6396. doi: 10.1002/ece3.2269

58. van Etten J. R package gdistance: Distances and routes on geographical grids. Journal of Statistical Software. 2017; 76(13):1–21. 10.18637/jss.v076.i13

59. Goslee SC, Urban DL. The ecodist package for dissimilarity-based analysis of ecological data. Journal of Statistical Software. 2007; 22:1–19. doi: 10.18637/jss.v022.i07

60. Ochoa-Ochoa LM, Mejía-Domínguez NR, Velasco JA, Marske KA, Rahbek C. Amphibian functional diversity is related to high annual precipitation and low precipitation seasonality in the New World. Global Ecology and Biogeography. 2019; 28(9):1219–1229. doi: 10.1111/geb.12926

61. Cohen MP, Alford RA. Factors affecting diurnal shelter use by the cane toad, *Bufo marinus*. Herpetologica. 1996; 52:72–181. https://www.jstor.org/stable/3892986

62. Aryal PC, Aryal C, Neupane S, Sharma B, Dhamala MK, Khadka D, et al. Soil moisture and roads influence the occurrence of frogs in Kathmandu Valley, Nepal. Global Ecology and Conservation. 2020; 23:e01197. doi: 10.1016/j.gecco.2020.e01197

63. Gould J, Beranek C, Valdez J, Mahony M. Quantity versus quality: A balance between egg and clutch size among Australian amphibians in relation to other life-history variables. Austral Ecology. 2022; 47(3):685–697. doi: 10.1111/aec.13154

64. Peterman WE. ResistanceGA: an R package for the optimization of resistance surfaces using genetic algorithms. Methods in Ecology and Evolution. 2018; 9:1638–1647. doi: 10.1111/2041-210X.12984

65. Peterman WE, Connette GM, Semlitsch RD, Eggert LS. Ecological resistance surfaces predict fine-scale genetic differentiation in a terrestrial woodland salamander. Molecular Ecology. 2014; 23(10):2402–2413. doi: 10.1111/mec.12747

66. Scrucca L. GA: A package for genetic algorithms in R. Journal of Statistical Software. 2013; 53:1–37. doi: 10.18637/jss.v053.i04

67. Bolker BM. Ecological Models and Data in R. Princeton: Princeton University Press; 2008.

68. Akaike H. A new look at the statistical model identification. IEEE Transactions on Automatic Control. 1974; 19(6):716–723. doi:10.1109/TAC.1974.1100705

69. Peterman WE, Pope NS. The use and misuse of regression models in landscape genetic analyses. Molecular Ecology. 2021; 30:37–47 doi: 10.1111/mec.15716

70. Luu K, Bazin E, Blum MG. pcadapt: an R package to perform genome scans for selection based on principal component analysis. Molecular Ecology Resources. 2017; 17(1):67–77. doi: 10.1111/1755-0998.12592

71. Storey JD, Tibshirani R. Statistical significance for genome wide studies. Proceedings of the National Academy of Sciences. 2003; 100:9440–9445, doi: 10.1073/pnas.1530509100

72. Forester BR, Lasky JR, Wagner HH, Urban DL. Comparing methods for detecting multilocus adaptation with multivariate genotype-environment associations. Molecular Ecology. 2018; 27(9):2215–2233. doi: 10.1111/mec.14584

73. Oksanen J, Blanchet FG, Kindt R, Legendre P, Minchin P, O’Hara RB, et al. vegan: Community Ecology Package. R package version 2.0-3. R Foundation for Statistical Computing, Vienna, Austria; 2012.

74. Frichot E, Schoville SD, Bouchard G, François O. Testing for associations between loci and environmental gradients using latent factor mixed models. Molecular Biology and Evolution. 2013; 30(7):1687–1699. doi: 10.1093/molbev/mst063

75. Frichot E, François O. LEA: An R package for landscape and ecological association studies. Methods in Ecology and Evolution. 2015; 6(8):925–929. doi: 10.1111/2041-210X.12382

76. de Villemereuil P, Frichot É, Bazin É, François O, Gaggiotti OE. Genome scan methods against more complex models: when and how much should we trust them? Molecular Ecology. 2014; 23(8):2006–2019. doi: 10.1111/mec.12705

77. Lotterhos KE, Whitlock MC. The relative power of genome scans to detect local adaptation depends on sampling design and statistical method. Molecular Ecology. 2015; 24:1031–1046. doi: 10.1111/mec.13100

78. Ge SX, Jung D, Yao R. ShinyGO: a graphical gene-set enrichment tool for animals and plants. Bioinformatics. 2020; 36(8):2628–2629. doi: 10.1093/bioinformatics/btz931

79. Ashburner M, Ball CA, Blake JA, Botstein D, Butler H, Cherry JM, et al. Gene ontology: tool for the unification of biology. Nature Genetics. 2000; 25(1):25–29. doi: 10.1038/75556

80. Kanehisa M, Furumichi M, Tanabe M, Sato Y, Morishima K. KEGG: new perspectives on genomes, pathways, diseases and drugs. Nucleic Acids Research. 2017; 45(D1):D353–D361. doi: 10.1093/nar/gkw1092

81. Allendorf FW. Genetics and the conservation of natural populations: allozymes to genomes. Molecular Ecology, 2017; 26:420–430. doi: 10.1111/mec.13948

82. Munshi-South J, Zak Y, Pehek E. Conservation genetics of extremely isolated urban populations of the northern dusky salamander (*Desmognathus fuscus*) in New York City. PeerJ. 2013; 1:e64. doi: 10.7717/peerj.64

83. Okamiya H, Kusano T. Lower genetic diversity and hatchability in amphibian populations isolated by urbanization. Population Ecology. 2018; 60(4):347–360. doi:10.1007/s10144-018-0627-4.

84. Wei X, Huang M, Yue Q, Ma S, Li B, Mu Z, et al. Long-term urbanization impacts the eastern golden frog (*Pelophylax plancyi*) in Shanghai City: Demographic history, genetic structure, and implications for amphibian conservation in intensively urbanizing environments. Evolutionary Applications. 2021; 14(1):117–135. doi: 10.1111/eva.13156

85. Arruda MP, Morielle-Versute E, Silva A, Schneider MPC, Gonçalves EC. Contemporary gene flow and weak genetic structuring in Rococo toad (*Rhinella schneideri)* populations in habitats fragmented by agricultural activities. Amphibia-Reptilia. 2011; 32(3):399–411. doi: 10.1163/017353711X588182

86. Stuart S, Hoffmann M, Chanson J, Cox N, Berridge R, Ramani P, Young B (eds). Threatened Amphibians of the World. Lynx Edicions, IUCN, and Conservation International, Barcelona, Spain; Gland, Switzerland; Arlington, Virginia; 2008.

87. McKee AM, Maerz JC, Smith LL, Glenn TC. Habitat predictors of genetic diversity for two sympatric wetland-breeding amphibian species. Ecology and Evolution. 2017; 7(16):6271–6283. doi: 10.1002/ece3.3203

88. Shaykevich DA, Pašukonis A, O’Connell LA. Long distance homing in the cane toad (*Rhinella marina*) in its native range. Journal of Experimental Biology. 2022; 225(2):jeb243048. doi: 10.1242/jeb.243048

89. Figueiredo-Vázquez C, Lourenço A, Velo-Antón G. Riverine barriers to gene flow in a salamander with both aquatic and terrestrial reproduction. Evolutionary Ecology. 2021; 35(3):483–511. doi: 10.1007/s10682-021-10114-z

90. Fonseca EM, Garda AA, Oliveira EF, Camurugi F, Magalhães FDM, Lanna FM, et al. The riverine thruway hypothesis: rivers as a key mediator of gene flow for the aquatic paradoxical frog Pseudis tocantins (Anura, Hylidae). Landscape Ecology. 2021; 36(10):3049–3060. doi: 10.1007/s10980-021-01257-z

91. Gutiérrez-Rodríguez J, Gonçalves J, Civantos E, Martínez-Solano I. Comparative landscape genetics of pond-breeding amphibians in Mediterranean temporal wetlands: The positive role of structural heterogeneity in promoting gene flow. Molecular Ecology. 2017; 26(20):5407–5420. doi: 10.1111/mec.14272

92. Gutiérrez-Rodríguez J, Gonçalves J, Civantos E, Maia-Carvalho B, Caballero-Díaz C, Gonçalves H, Martínez-Solano Í. The role of habitat features in patterns of population connectivity of two Mediterranean amphibians in arid landscapes of central Iberia. Landscape Ecology. 2023; 38(1):99–116. doi: 10.1007/s10980-022-01548-z

93. Albero L, Martínez-Solano Í, Hermida M, Vera M, Tarroso P, Bécares E. Open areas associated with traditional agriculture promote functional connectivity among amphibian demes in Mediterranean agrosystems. Landscape Ecology. 2023; 38(12):3045–3059. doi: 10.1007/s10980-023-01725-8

94. Fusco NA, Pehek E, Munshi-South J. Urbanization reduces gene flow but not genetic diversity of stream salamander populations in the New York City metropolitan area. Evolutionary Applications. 2021; 14(1):99–116. doi: 10.1111%2Feva.13025

95. Parsley MB, Torres ML, Banerjee SM, Tobias ZJ, Goldberg CS, Murphy MA, Mims MC. Multiple lines of genetic inquiry reveal effects of local and landscape factors on an amphibian metapopulation. Landscape Ecology. 2020; 35:319–335. doi: 10.1007/s10980-019-00948-y

96. Spear SF, Storfer A. Anthropogenic and natural disturbance lead to differing patterns of gene flow in the Rocky Mountain tailed frog, *Ascaphus montanus*. Biological Conservation. 2010; 143(3):778–786. doi: 10.1016/j.biocon.2009.12.021

97. Spear SF, Storfer A. Landscape genetic structure of coastal tailed frogs (*Ascaphus truei*) in protected vs. managed forests. Molecular Ecology. 2008; 17(21):4642–4656. doi: 10.1111/j.1365-294X.2008.03952.x

98. Bray SJ. Notch signaling in context. Nature Reviews Molecular Cell Biology. 2016; 17(11):722–735. doi: 10.1038/nrm.2016.94

99. Zhou B, Lin W, Long Y, Yang Y, Zhang H, Wu K, Chu Q. Notch signaling pathway: architecture, disease, and therapeutics. Signal Transduction and Targeted Therapy. 2022; 7(1):95. doi: 10.1038/s41392-022-00934-y

100. Coffman C, Harris W, Kintner C. Xotch, the *Xenopus* homolog of *Drosophila* notch. Science. 1990; 249(4975);1438–1441. doi: 10.1126/science.2402639

101. Favarolo MB, López SL. Notch signaling in the division of germ layers in bilaterian embryos. Mechanisms of Development. 2018; 154:122–144. doi: 10.1016/j.mod.2018.06.005

102. Krens SG, Spaink HP, Snaar-Jagalska BE. Functions of the MAPK family in vertebrate- development. FEBS Letters. 2006; 580(21):4984–4990. doi: 10.1016/j.febslet.2006.08.025

103. Wen X, Jiao L, Tan H. MAPK/ERK pathway as a central regulator in vertebrate organ regeneration. International Journal of Molecular Sciences. 2022; 23(3):1464. doi: 10.3390/ijms23031464

104. Haigh JJ. Role of VEGF in organogenesis. Organogenesis. 2008; 4(4):247–256. doi: 10.4161/org.4.4.7415

105. Shibuya M. Vascular endothelial growth factor and its receptor system: physiological functions in angiogenesis and pathological roles in various diseases. The Journal of Biochemistry. 2013; 153(1):13–19. doi: 10.1093/jb/mvs136

106. Vieira JM, Ruhrberg C, Schwarz Q. VEGF receptor signaling in vertebrate development. Organogenesis. 2010; 6(2):97–106. doi: 10.4161/org.6.2.11686

107. Gupta A, Storey KB. A “notch” in the cellular communication network in response to anoxia by wood frog (*Rana sylvatica*). Cellular Signalling. 2022; 93:110305. doi: 10.1016/j.cellsig.2022.110305

108. Wells KD. The ecology and behavior of amphibians. Chicago: University of Chicago Press; 2019.

109. Sanuy D, Oromí N, Galofré A. Effects of temperature on embryonic and larval development and growth in the natterjack toad (*Bufo calamita*) in a semi-arid zone. Animal Biodiversity and Conservation. 2008; 31(1):41–46. 10.32800/abc.2008.31.0041

110. Chen X, Ren C, Teng Y, Shen Y, Wu M, Xiao H, Wang H. Effects of temperature on growth, development and the leptin signaling pathway of *Bufo gargarizans*. Journal of Thermal Biology. 2021; 96:102822. doi: 10.1016/j.jtherbio.2020.102822

111. Lorrain-Soligon L, Bizon T, Robin F, Jankovic M, Brischoux F. Variations of salinity during reproduction and development affect ontogenetic trajectories in a coastal amphibian. Environmental Science and Pollution Research. 2024; 31:11735–11748. doi: 10.1007/s11356-024-31886-1

112. Maciel TA, Juncá FA. Effects of temperature and volume of water on the growth and development of tadpoles of *Pleurodema diplolister* and *Rhinella granulosa* (Amphibia: Anura). Zoologia (Curitiba). 2009; 26:413–418. doi: 10.1590/S1984-46702009000300005

113. Wijethunga U, Greenlees M, Shine R. Moving south: effects of water temperatures on the larval development of invasive cane toads (*Rhinella marina*) in cool-temperate Australia. Ecology and Evolution. 2016; 6(19):6993–7003. doi: 10.1002/ece3.2405

114. Brans KI, Engelen JM, Souffreau C, De Meester L. Urban hot-tubs: Local urbanization has profound effects on average and extreme temperatures in ponds. Landscape and Urban Planning. 2018; 176:22–29. doi: 10.1016/j.landurbplan.2018.03.013

115. Wang S, Kuang Y, Liang L, Sun B, Zhao X, Zhang L, Chang Y. Resequencing and SNP discovery of Amur ide (*Leuciscus waleckii*) provides insights into local adaptations to extreme environments. Scientific Reports. 2021; 11(1):5064. doi: 10.1038/s41598-021-84652-5

116. Mandinova, A, Lefort K, Di Vignano AT, Stonely W, Ostano P, Chiorino G, et al. The FoxO3a gene is a key negative target of canonical Notch signalling in the keratinocyte UVB response. The EMBO Journal. 2008; 27(8):1243–1254. doi: 10.1038/emboj.2008.45

117. Zhang XZ, Ma XD, Wang WT, Peng F, Hou YM, Shen YX, et al. Comparative skin histological and transcriptomic analysis of *Rana kukunoris* with two different skin colors. Comparative Biochemistry and Physiology Part D: Genomics and Proteomics. 2024; 50:101217. doi: 10.1016/j.cbd.2024.101217

118. Blaustein AR, Belden LK. Amphibian defenses against ultraviolet-B radiation. Evolution & Development. 2003; 5(1):89–97. doi: 10.1046/j.1525-142X.2003.03014.x

119. Kosch TA, Torres-Sánchez M, Liedtke HC, Summers K, Yun MH, Crawford AJ, et al. The Amphibian Genomics Consortium: advancing genomic and genetic resources for amphibian research and conservation. bioRxiv 2024.06.27.601086 [Preprint]. 2024 Available from: doi: 10.1101/2024.06.27.601086

120. Franco-Belussi L, De Oliveira C. The spleen of *Physalaemus nattereri* (Amphibia: Anura): morphology, melanomacrophage pigment compounds and responses to α-melanocyte stimulating hormone. Italian Journal of Zoology. 2016; 83(3):298–305. doi: 10.1080/11250003.2016.1194488

121. Zhou J, Liao Z, Liu Z, Guo X, Zhang W, Chen Y. Urbanization increases stochasticity and reduces the ecological stability of microbial communities in amphibian hosts. Frontiers in Microbiology. 2023; 13:1108662. doi: 10.3389/fmicb.2022.1108662

122. Grogan LF, Robert J, Berger L, Skerratt LF, Scheele BC. Castley JG, et al. Review of the amphibian immune response to chytridiomycosis, and future directions. Frontiers in Immunology. 2018; 9:2536. doi: 10.3389/fimmu.2018.02536

123. Huang L, Li J, Anboukaria H, Luo Z, Zhao M, Wu H. Comparative transcriptome analyses of seven anurans reveal functions and adaptations of amphibian skin. Scientific Reports. 2016; 6(1):24069. doi: 10.1038/srep24069

124. Varga JF, Bui-Marinos MP, Katzenback BA. Frog skin innate immune defenses: sensing and surviving pathogens. Frontiers in Immunology. 2019; 9:433806. doi: 10.3389/fimmu.2018.03128

125. Chuphal B, Rai U, Roy B. Teleost NOD-like receptors and their downstream signaling pathways: A brief review. Fish and Shellfish Immunology Reports. 2022; 3:100056. doi: 10.1016/j.fsirep.2022.100056

126. Geijtenbeek TB, Gringhuis SI. Signaling through C-type lectin receptors: shaping immune responses. Nature Reviews Immunology. 2009; 9(7):465–479. doi: 10.1038/nri2569

127. Nie L, Cai SY, Shao JZ, Chen J. Toll-like receptors, associated biological roles, and signaling networks in non-mammals. Frontiers in Immunology. 2018; 9:391632. doi: 10.3389/fimmu.2018.01523

128. Rehwinkel J, Gack MU. RIG-I-like receptors: their regulation and roles in RNA sensing. Nature Reviews Immunology. 2020; 20(9):537–551. doi: 10.1038/s41577-020-0288-3

129. Garcia-Vidal C, Carratalà J. Patogenia de la infección fúngica invasora. Enfermedades Infecciosas y Microbiología Clínica. 2012; 30(3):151–158. doi: 10.1016/j.eimc.2011.09.011

130. Jiang N, Fan Y, Zhou Y, Meng Y, Liu W, Li Y, et al. The immune system and the antiviral responses in Chinese giant salamander, *Andrias davidianus*. Frontiers in Immunology. 2021; 12:718627. doi: 10.3389/fimmu.2021.718627

131. Ceccato E, Cramp RL, Seebacher F, Franklin CE. Early exposure to ultraviolet-B radiation decreases immune function later in life. Conservation Physiology. 2016; 4(1):cow037. doi: 10.1093/conphys/cow037

132. Yang W, Qi Y, Fu J. Genetic signals of high-altitude adaptation in amphibians: a comparative transcriptome analysis. BMC Genetics. 2016; 17:1–10. doi: 10.1186/s12863-016-0440-z

133. McDonald CA, Becker CG, Lambertini C, Toledo LF, Haddad CF, Zamudio KR. Host immune responses to enzootic and invasive pathogen lineages vary in magnitude, timing, and efficacy. Molecular Ecology. 2023; 32(9):2252–2270. doi: 10.1111/mec.16890

134. Rödin-Mörch P, Palejowski H, Cortazar-Chinarro M, Kärvemo S, Richter-Boix A, Höglund J, Laurila A. Small-scale population divergence is driven by local larval environment in a temperate amphibian. Heredity. 2021; 126(2):279–292. doi: 10.1038/s41437-020-00371-z

135. Brown GP, Phillips BL, Dubey S, Shine R. Invader immunology: invasion history alters immune system function in cane toads (*Rhinella marina*) in tropical Australia. Ecology Letters. 2015; 18(1):57–65. doi: 10.1111/ele.12390

136. Rollins LA, Richardson MF, Shine R. A genetic perspective on rapid evolution in cane toads (*Rhinella marina*). Molecular Ecology. 2015; 24:313–327. doi: 10.1111/mec.13184

137. Medina R, Wogan GOU, Bi K, Termignoni-García F, Bernal MH, Jaramillo-Correa JP, Wang IJ, Vázquez-Domínguez E. Phenotypic and genomic diversification with isolation by environment along elevational gradients in a neotropical treefrog. Molecular Ecology. 2021; 30(16):4062–4076. doi: 10.1111/mec.16035

138. Goessens T, De Baere S, Deknock A, De Troyer N, Van Leeuwenberg R, Martel A, et al. Agricultural contaminants in amphibian breeding ponds: Occurrence, risk and correlation with agricultural land use. Science of the Total Environment. 2022; 806:150661. doi: 10.1016/j.scitotenv.2021.150661

139. Jiménez RR, Sommer S. The amphibian microbiome: natural range of variation, pathogenic dysbiosis, and role in conservation. Biodiversity and Conservation. 2017; 26:763–786. doi: 10.1007/s10531-016-1272-x

140. Preuss JF, Greenspan SE, Rossi EM, Lucas Gonsales EM, Neely WJ, Valiati VH, et al. Widespread pig farming practice linked to shifts in skin microbiomes and disease in pond- breeding amphibians. Environmental Science & Technology. 2020; 54(18):11301–11312. doi: 10.1021/acs.est.0c03219

141. Meng Y, Fan Y, Zhou Y, Jiang N, Xue M, Liu W, et al. Identification and comparative expression analysis of RIG-I and MDA5 in Chinese giant salamander *Andrias davidianus*. Aquaculture Research. 2020; 51(11):4575–4582. doi: 10.1111/are.14803

142. Li F, Chen B, Xu M, Feng Y, Deng Y, Huang X, et al. Immune activation and inflammatory response mediated by the nod/toll-like receptor signaling pathway—The potential mechanism of Bullfrog (*Lithobates catesbeiana*) meningitis caused by *Elizabethkingia miricola*. International Journal of Molecular Sciences. 2023; 24(19):14554. doi: 10.3390/ijms241914554

